# An endothelial SOX18-mevalonate pathway axis enables repurposing of statins for infantile hemangioma

**DOI:** 10.1101/2024.01.29.577829

**Authors:** Annegret Holm, Matthew S. Graus, Jill Wylie-Sears, Luke Borgelt, Jerry Wei Heng Tan, Sana Nasim, Long Chung, Ashish Jain, Mingwei Sun, Liang Sun, Pascal Brouillard, Ramrada Lekwuttikarn, Harry Kozakewich, Jacob Yanfei Qi, Joyce C. Teng, John B. Mulliken, Miikka Vikkula, Mathias Francois, Joyce Bischoff

## Abstract

Infantile hemangioma (IH) is the most common tumor in children and a paradigm for pathological vasculogenesis, angiogenesis and regression. Propranolol is the mainstay of treatment for IH. It inhibits hemangioma vessel formation via a β-adrenergic receptor independent off-target effect of its R(+) enantiomer on the endothelial specific transcription factor sex-determining region Y (SRY) box transcription factor 18 (SOX18). Transcriptomic profiling of patient-derived hemangioma stem cells uncovered the mevalonate pathway (MVP) as a target of R(+) propranolol. Loss of SOX18 function confirmed R(+) propranolol mode of action on the MVP. Functional validation in preclinical IH models revealed that statins - targeting the MVP - are potent inhibitors of hemangioma vessel formation. We propose a novel SOX18-MVP-axis as a central regulator of IH pathogenesis and suggest statin repurposing to treat IH. Our findings reveal novel pleiotropic effects of beta-blockers and statins acting on the SOX18-MVP axis to disable an endothelial specific program in IH, which may impact other scenarios involving pathological vasculogenesis and angiogenesis.

**Graphical abstract:** 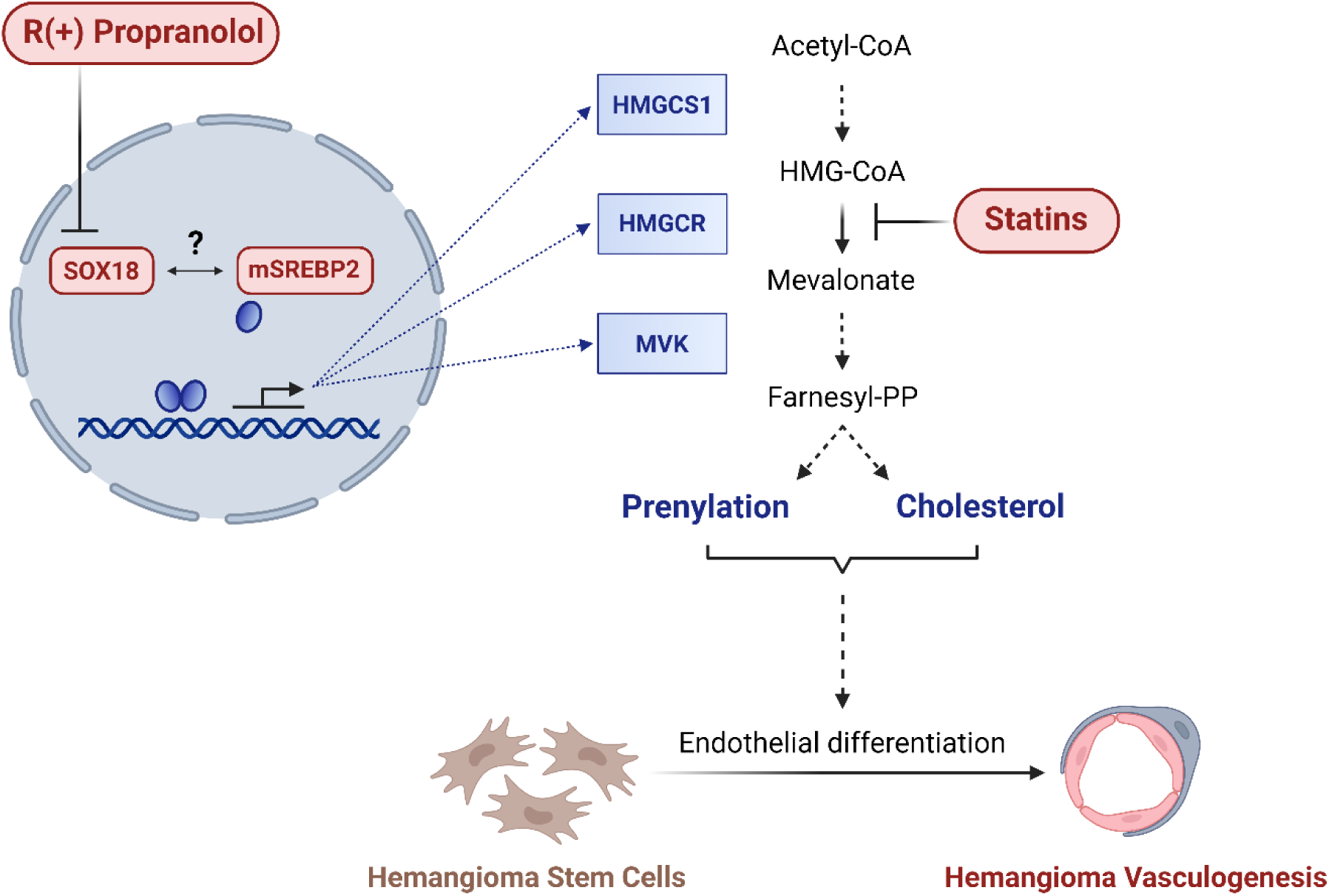

## INTRODUCTION

Infantile hemangioma (IH) is a common benign vascular tumor of infancy with an incidence of 2-10%. It predominantly occurs in female and premature infants of European descent. IH arises postnatally at 3-7 weeks of age and grows rapidly during the proliferating phase, which can continue for 12 months on average. A spontaneous and gradual involuting phase follows spanning 2-7 years. A subtype of IH - often involving the oral mucosa - do not undergo complete involution and may regrow after treatment discontinuation. For most children, IH poses no serious risk; however, 10-15% of lesions require treatment to prevent sequelae such as disfigurement and functional impairment including vision loss and airway obstruction, consumptive hypothyroidism, and high-output cardiac failure (1). Propranolol was discovered serendipitously to be effective for IH. It is currently the only FDA-approved drug for IH, based on a randomized controlled clinical trial (2). Despite its successful repurposing for IH, propranolol is associated with beta-adrenergic side effects including hypotension, bradycardia, peripheral vasospasm, hypoglycemia and seizures, bronchospasm, slowed weight gain, diarrhea, agitation, sleep disturbance, and potentially adverse neurocognitive outcomes (2–5). The complete response rate to propranolol is reported to be 60%; a subset of IH regrow after treatment is discontinued (2). These safety and efficacy concerns underscore the need for additional treatment options for patients with IH.

It is firmly established that propranolol acts as a non-selective antagonist of the G protein-coupled β1- and β2-adrenergic receptors (GPCR). Propranolol consists of an equimolar (1:1) mixture of S(-) and R(+) enantiomers. The S(-) enantiomer is a potent antagonist of β1 and β2-adrenergic receptors, while the R(+) enantiomer is largely devoid of beta-blocker activity (6). This provided an opportunity to identify a novel R(+) propranolol-dependent pathway in hemangioma stem cells (HemSC). HemSC are isolated from proliferating phase patient IH specimens and recapitulate hemangiogenesis in nude mice (7). R(+) propranolol inhibits HemSC endothelial differentiation in vitro and HemSC vasculogenesis in a pre-clinical model (8, 9). Furthermore, we established that R(+) propranolol interferes with the activity of the transcription factor (TF) SRY (sex-determining region Y) HMG box-containing 18 (SOX18).

Pharmacological interference has also been observed with Sm4 - a small molecule inhibitor of SOX18 - further validating the role of this molecular target in IH (8, 9). These findings have led to the repurposing of propranolol in patients with hypotrichosis-lymphedema-telangiectasia-renal defect syndrome (HLTRS) - a rare vascular disease caused by a dominant-negative truncating mutation in SOX18 (10). Together, these observations provided evidence for the pharmaco-genetic interaction between SOX18 and propranolol in human disease (8). In summary, we have identified a SOX18-dependent inhibition of HemSC endothelial differentiation and vessel formation in vivo as the molecular basis of propranolol-mediated inhibition of IH.

SOX18 is a master regulator of vascular development and endothelial specification and is expressed in nascent differentiating blood and lymphatic endothelium as well as in endothelial progenitor cells in adults (11). It plays fundamental roles in arterial specification (12), lymphangiogenesis (13) and tumor angiogenesis (14). Because of its known role during embryogenesis, this prompted us to investigate the molecular basis of its pharmacological blockade in the context of HemSC endothelial differentiation.

To elucidate the functional role of SOX18 in IH vasculogenesis, we set out to identify genes whose expression in differentiating HemSC is altered by R(+) propranolol. We discovered that R(+) propranolol coordinately downregulates transcripts encoding enzymes in the mevalonate pathway (MVP). The MVP is central to cholesterol and isoprenoid biosynthesis and is controlled by the rate-limiting enzyme 3-hydroxy-3-methylglutaryl-CoA reductase A (HMGCR) (15), which produces mevalonate. Additional regulators are the transcription factor sterol regulatory element binding protein-2 (SREBP2) that induces expression of MVP genes (16) and the ATP-binding cassette transporter A1 (ABCA1) that acts as a negative regulator of SREBP2 by facilitating retrograde transport of cholesterol from the plasma membrane to the endoplasmic reticulum (ER) (17). Statins, competitive HMGCR inhibitors, are widely prescribed to reduce low-density-lipoprotein cholesterol in patients at risk for cardiovascular disease (18). Pleiotropic activities of statins beyond the lipid-lowering effect have been established (19–21). Accumulating evidence suggests a role for statins in epigenetic modifications in a cardiovascular disease and an oncology context (22–24). Here, we uncover a novel molecular relationship between an endothelial specific TF and the MVP. We show that R(+) propranolol causes a SOX18-mediated downregulation of MVP genes in HemSC during their endothelial differentiation. Blocking the MVP with statins inhibits endothelial differentiation and vessel formation, suggesting that statins could be repurposed to safely treat the vascular overgrowth in IH.

## RESULTS

### R(+) propranolol globally reduces transcript levels of MVP genes in patient-derived HemSC undergoing endothelial differentiation

To identify the downstream targets of R(+) propranolol in HemSC to hemangioma endothelial cell (HemEC) differentiation in a non-biased way, we performed bulk RNA sequencing of HemSC isolated from 6 different IH specimens. Table 1 provides an overview of patient samples used in respective experiments. The HemSC were induced to undergo endothelial differentiation for 6 days and then treated with or without R(+) propranolol (20uM) for 2 hours (Supplemental Figure 1.1). The timing and dosage were determined by previously observed downregulation of *NOTCH1* expression, a widely known *SOX18* transcriptional target, under this treatment condition (9). HemSC at Day 4 of endothelial differentiation, prior to onset of endothelial marker expression, were analyzed as well. The Kyoto Encyclopedia of Genes and Genomes (KEGG) analysis identified steroid biosynthesis as differentially affected at Day 6 (Supplemental Figure 1.2), which was confirmed by gene ontology analysis of the subontology *biological processes* (8 of the top 20 processes are related to cholesterol biosynthesis) (Figure 1A). Subsequent gene by gene analysis revealed R(+) propranolol treatment on Day 6 significantly reduced transcripts encoding several enzymes of the MVP, including the rate-limiting enzyme HMGCR as well as 3-hydroxy-3-methylglutaryl-CoA synthase 1 (HMGCS1) and mevalonate kinase (MVK), while few changes were seen at Day 4 (Figure 1B,C; Supplemental Table S1). The significant increase in *SOX18* mRNA from Day 4 to Day 6 of differentiation is consistent with the onset of MVP gene sensitivity to R(+) propranolol at Day 6 (Figure 1D). Downregulation of *HMGCS1*, *HMGCR*, and *MVK* as well as upregulation of *ABCA1 -* a negative regulator of SREBP2 transcriptional activity - was confirmed by qPCR in cells treated with R(+) propranolol for 2 hours or for 4 days of the differentiation protocol (Figure 1E). Overall, 105 genes in HemSC were differentially expressed at Day 4 versus 2482 genes at Day 6 of differentiation. Differential expression was defined as log_2_ fold change >1 and adjusted p-value <0.05. The increase in differentially expressed genes at Day 6 defines a critical time window for *SOX18* inhibition and therefore R(+) propranolol effect on HemSC differentiation. This concept is further supported by the lack of downregulation of MVP transcripts seen in HemSC at Day 4 of differentiation, prior to endothelial marker expression. RNA-Seq data will be submitted to the Gene Expression Omnibus archive at the National Center for Biotechnology Information.

**Figure 1.**
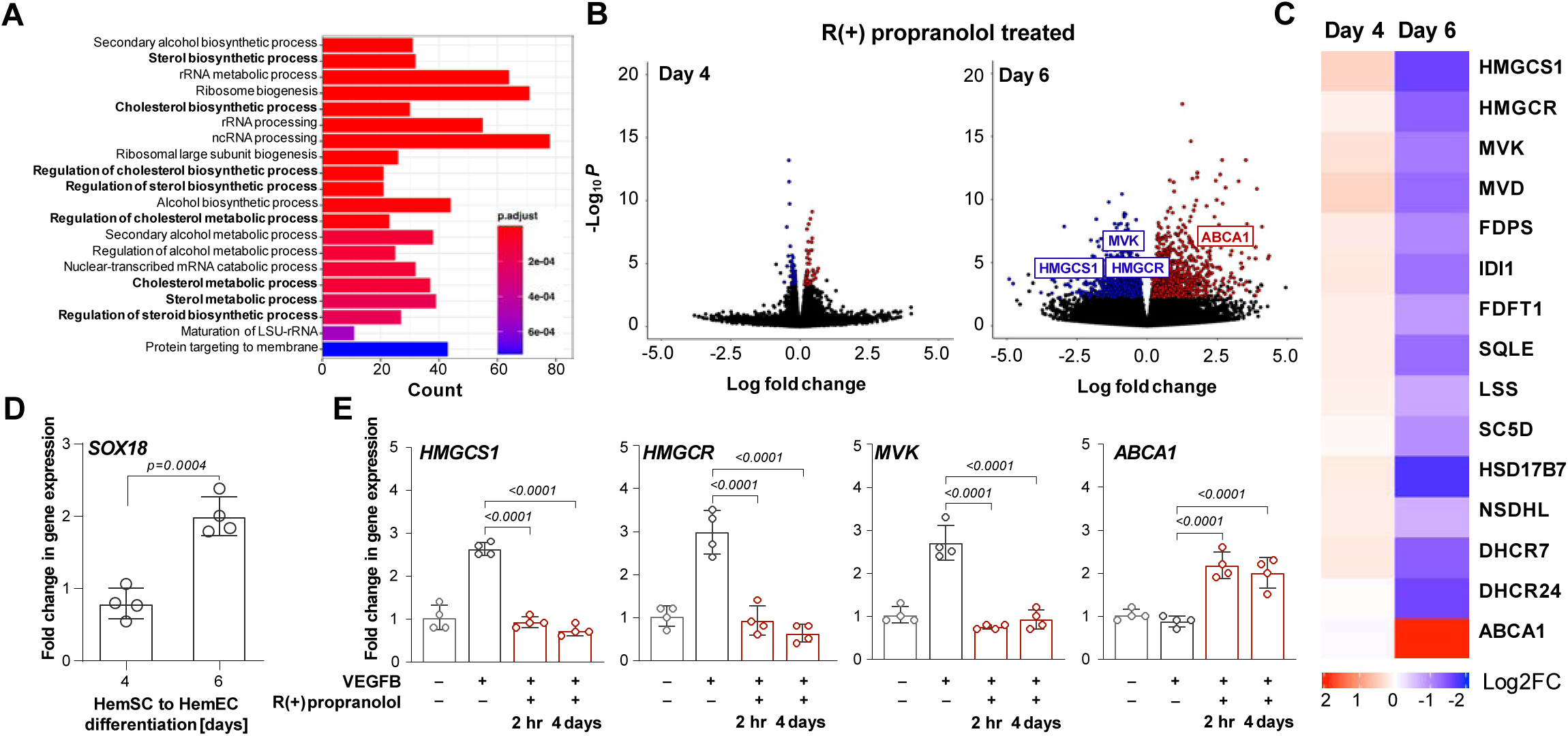
R(+) propranolol reduces MVP transcripts in IH -derived HemSC undergoing endothelial differentiation. (**A**) Gene ontology analysis of the subontology *biological processes* of bulk RNA-seq data set from IH-derived HemSC (n=6). (**B**) Volcano plots show an increase in differentially expressed genes on Day 6 compared to Day 4 of HemSC to endothelial differentiation. (**C**) Heat map with MVP gene transcripts at Day 4 and Day 6 as well as ABCA1 – a negative regulator of the MVP transcription factor SREBP2. (**D**) SOX18 measured by qPCR is significantly increased on Day 6 compared to Day 4 (n=3). (**E**) HemSC induced to undergo endothelial differentiation were treated with R+ propranolol for 2 hours on Day 6, or from Day 2-6, i.e., 4-day treatment. qPCR analyses for three MVP genes - *HMGCS1*, *HMGCR*, *MVK,* as well as *ABCA1,* a negative regulator of the MVP transcription factor SREBP2 (n=4). P values were calculated using a 2- tailed, unpaired t-test (**D**) and by a one-way ANOVA multiple comparisons test with Šidák-correction (**E**). Data show the mean ± SD. Data represent sample sizes n=4 (**D**,**E**).

**Table 1:** Overview of patient cells and tissues used throughout the study.

| Sample ID | Gender | Age at resection | Figure |
| --- | --- | --- | --- |
| ***Proliferating IH (<1 year)*** |  |  |  |
| P-150 | M | 3m | 1, 3, 4, 5 |
| P-147 | M | 3m | 4, 5 |
| P-138 | F | 3m | 4 |
| P-133 | M | 3m | 4, 5 |
| P-171 | F | 4m | 1, 3, 4, 5 |
| P-165 | F | 4m | 1, 3, 5 |
| P-168 | F | 5m | 4 |
| P-125 | F | 5m | 1, 4 |
| P-167 | F | 7m | 1, 4 |
| P-149 | F | 7m | 1, 3, 4, 5 |
| ***Involuting IH (>1 year)*** |  |  |  |
| I-69 | F | 1y 1m | 4 |
| I-79 | F | 1y 6m | 4 |
| I-71 | F | 2y 1m | 4 |
| I-85 | M | 2y 2m | 4 |
| I-99 | M | 2y 6m | 4 |
| I-84 | M | 2y 6m | 4 |
| I-78 | M | 2y 6m | 4 |
| I-67 | M | 2y 6m | 4 |
| I-81 | F | 2y 11m | 4 |
| I-68 | F | 2y 11m | 4 |
| ***Rebounding IH*** |  |  |  |
| R-1 | F | 6y 4m | 4 |
| R-2 | F | 9y 6m | 4 |
| ***Skin Controls*** |  |  |  |
| C-1 | F | 8m | 4 |
| C-2 | M | 1y | 4 |
| C-3 | M | 3y 7m | 4 |
| C-4 | M | 5y 4m | 4 |

To further elucidate the effect of SOX18 inhibition on the MVP, we investigated SOX18-binding locations in the genome using a publicly available ChIP-Seq dataset in human umbilical endothelial cells (HUVEC) (25). We discovered SOX18 binding locations within the *HMGCS1* and *HMGCR* gene loci that corresponded with ENCODE candidate cis-regulatory elements (Figure 2A). The RNA-Seq experiments, combined with the identification of SOX18-binding sites in regulatory regions of *HMGCS1* and *HMGCR*, support the ability of R(+) propranolol to interfere with SOX18 directed transcriptional regulation of critical genes in or related to the MVP.

**Figure 2.**
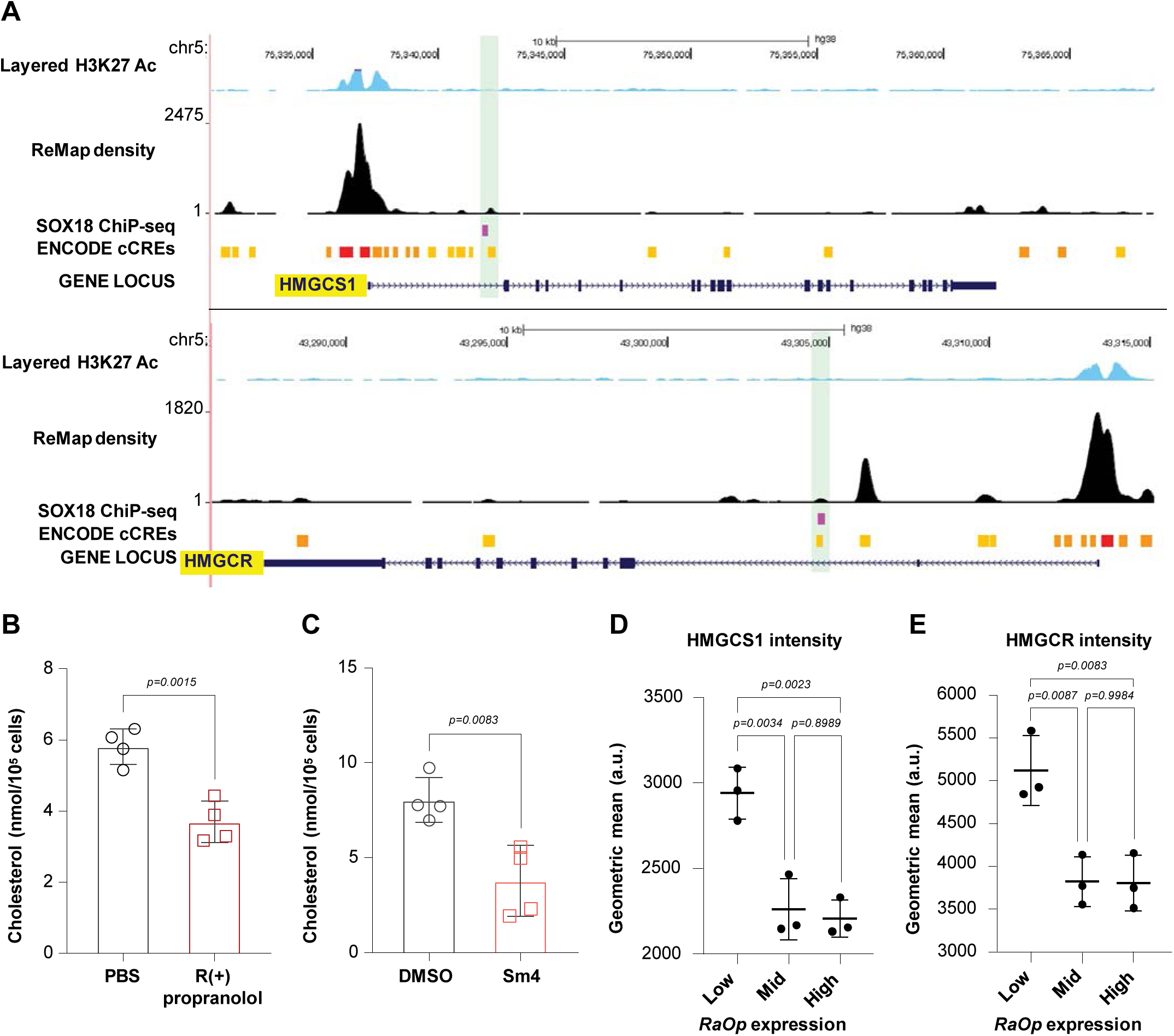
SOX18 regulates the MVP in endothelial cells. (**A**) ChIP-Seq dataset in HUVEC demonstrates SOX18 binding sites within the *HMGCS1* and *HMGCR* gene loci which correspond with ENCODE candidate cis-regulatory elements. (**B**,**C**) A functional role for SOX18 in cholesterol biosynthesis was tested by pharmacological disruption of SOX18 activity with R+ propranolol or the SOX18 inhibitor Sm4. HUVECs were cholesterol depleted by incubation with methyl beta-cyclodextrin (MBCD) for 16 hours, followed by treatment ± R(+) propranolol or ± Sm4 for 16 hours each. Endogenous cholesterol levels were measured by mass spectrometry. (**D**,**E**) Overexpression of Ragged Opossum (*RaOp*), a dominant negative version of SOX18 that is known to disrupt SOX18 activity, decreased immunofluorescent staining of HMGCS1 and HMGCR in HUVECs. P values were calculated using a 2- tailed, unpaired t-test (**B**,**C**) and by a one-way ANOVA multiple comparisons test with Tukey-correction (**D**,**E**). Data show the mean ± SD. Data represent sample sizes n=4 (**B**,**C**) and n=3 (**D**,**E**).

### SOX18 activity positively regulates MVP biosynthetic output in human endothelial cells

To validate the role of SOX18 in regulating genes in the MVP, we tested whether pharmacological disruption of SOX18 activity through treatment with R(+) propranolol or the known SOX18 inhibitor Sm4 (26) would have an effect on cholesterol biosynthesis in HUVEC. HUVEC were first cholesterol-depleted by incubation with methyl beta-cyclodextrin (MBCD) and treated ±R(+) propranolol or ± Sm4 for 16 hours, allowing for new cholesterol synthesis, which was then measured by targeted mass spectrometry. Endogenous cholesterol levels were significantly reduced in both R(+) propranolol- and Sm4-treated HUVEC, consistent with SOX18-mediated downregulation of MVP gene expression (Figure 2B,C). Conversely, we tested the effect of an overexpressed, dominant negative version of SOX18 (*Ragged Opossum* (*RaOp*)) that is known to disrupt SOX18 activity (27). High expression of SOX18*^RaOp^*decreased immunofluorescent staining for HMGCS1 and HMGCR compared to low SOX18*^RaOp^* indicating a gene dose response effect on two key MVP enzymes (Figure 2D,E; Supplemental Figure 2.1, 2.2). Taken together, both experiments demonstrate that SOX18 activity positively regulates cholesterol biosynthesis in an endothelial context.

### R(+) propranolol mode of action on the MVP is mediated via a SOX18-/SREBP2-dependent mechanism

SREBP2 is the master transcriptional regulator of the MVP and is ubiquitously expressed. We postulate that in HemSC undergoing endothelial differentiation, SREBP2 and SOX18 interact to co-regulate the MVP. To test this idea, we analyzed the effect of R(+) propranolol on levels of the SREBP2 precursor (inactive), which is anchored in the endoplasmic reticulum, and the 62kDa NH2 basic helix loop helix (bHLH) leucine zipper domain (16, 28). The 125 kDa precursor is proteolytically cleaved to release the 62kDa mature form when cholesterol levels drop. The mature 62kDa form of SREBP2 then translocates into the nucleus to activate transcription of MVP genes. We hypothesized that R(+) propranolol disruption of SOX18 would reduce 62 kDa SREBP2 in HemSC undergoing endothelial differentiation.

HemSC (n=4) were treated for 2 hours ± R(+) propranolol on Day 6 of VEGF-B induced endothelial differentiation. Cell lysates analyzed by Western Blot (WB) showed that R(+) propranolol decreased mature 62 kDa SREBP2 (Figure 3A,B). Rapid turnover of mature SREBP2 within 4 hours has been previously demonstrated (29). The reduced levels of 62kDa SREBP2 upon R(+) propranolol treatment are consistent with the transcriptomic findings from bulk RNA-seq and qPCR and suggest SOX18 and SREBP2 interact to regulate the MVP in nascent HemEC.

**Figure 3.**
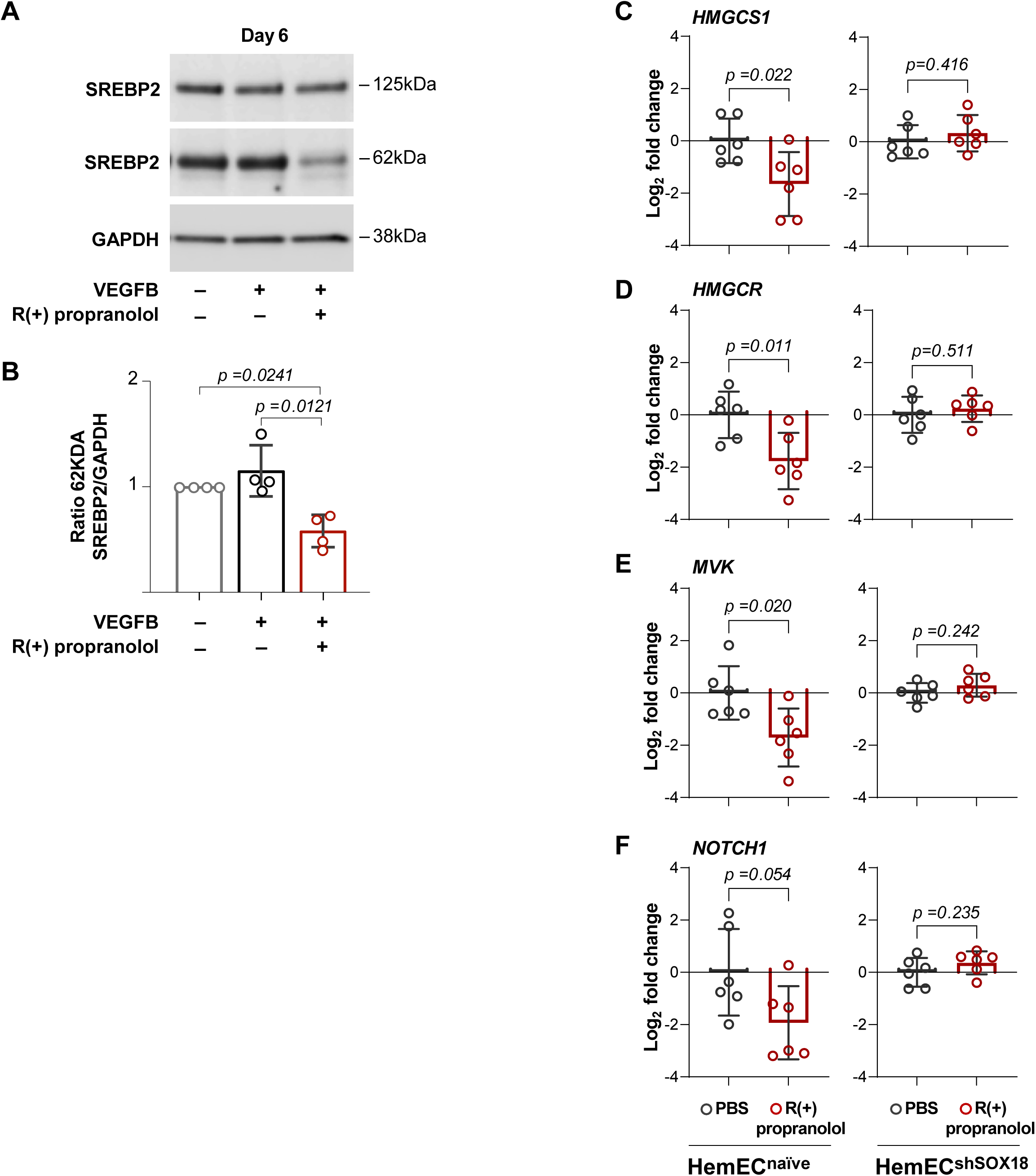
R(+) propranolol mode of action on the MVP is mediated via a SOX18-/SREBP2-dependent mechanism. (**A**,**B**) Patient-derived HemSC undergoing endothelial differentiation were treated for 2 hours ± R(+) propranolol on Day 6 and analyzed by Western blot with anti-SREBP2 (n=4). ImageJ quantification shows decreased levels of 62kDa mature SREBP2 in R(+) propranolol treated cells. (**C- F**) Lentiviral knockdown of SOX18 in patient-derived HemEC (n=3). Log_2_ fold changes in *HMGCS1*, *HMGCR* and *MVK* show significant reduction upon R(+) propranolol treatment (red) in naïve HemEC, while this effect is not evident in HemEC^shSOX18^; data normalized to PBS treated cells (black). *NOTCH1*, a known SOX18 target gene, served as a positive control. P values were calculated using one-way ANOVA multiple comparisons test with Šidák-correction (**B**) and by a 2-tailed, unpaired t-test (**C**-**F**). Data show the mean ± SD. Data represent sample sizes for n=4 (**B**) and n=6 naïve HemEC (denoted as HemEC 133,150,171) and n=6 HemEC^shSOX18^ with knockdown of SOX18 performed twice in each HemEC line (efficiency >70%) in (**C**-**F**).

We next assessed SOX18-dependency of the R(+) propranolol mediated effects by genetic knockdown of SOX18 using shRNA. We used differentiated IH-derived HemEC for this experiment because HemEC express high levels of SOX18. Lentiviral knockdown of SOX18 in three different IH-derived HemEC was conducted and the extent of SOX18 knockdown was verified by qPCR and WB (Supplemental Figure 3.1,2). HemEC with a SOX18 knockdown efficiency >70% were used. The requirement for SOX18 in R(+)- mediated downregulation of MVP genes was analyzed by qPCR measurement of *HMGCS1* and *HMGCR* and *MVK* in naïve HemEC and HemEC^shSOX18^. R(+) propranolol did not reduce the mRNA levels of MVP genes in HemEC^shSOX18^ (Figure 3C-E). The known SOX18 target *NOTCH1* served as a positive control for R(+) propranolol (Figure 3F). In summary, the knockdown of SOX18 in HemEC abolished the R(+) propranolol effect on mRNA transcript levels of *HMGCS1*, *HMGCR* and *MVK*, showing via genetic disruption of SOX18 that there is a transcriptional link between SOX18 and the MVP in IH derived cells (Figure 1-3).

### Nuclear SOX18 and SREBP2 in proliferating phase and regrowing IH indicate active MVP

Based on our transcriptomic and biochemical characterization from patient-derived HemSC and HemEC, we investigated the expression and localization of SOX18 and SREBP2 in patient IH histological sections of different tumor phases. We stained proliferating and involuting IH tissue sections (n=10, each) from specimens excised from children with IH who had not been treated with propranolol or corticosteroid (or any other treatment). Immunofluorescent staining with antibodies against human SOX18 and SREBP2, and with the lectin *Ulex Europaeus Agglutinin I* (UEA1), which specifically binds to human endothelial cells, showed co-localization of SOX18 and SREBP2 in endothelial nuclei throughout proliferating phase IH sections. Age-matched human skin (n=4), stained in parallel for comparison, was devoid of SOX18. Furthermore, SOX18 was largely absent from vessels in involuting phase IH. SOX18 is not routinely detected in mature, quiescent blood vessels, as it is not needed to maintain an endothelial phenotype (30). We next quantified SOX18^+^SREBP2^+^ nuclei/endothelial nuclei as a metric for active MVP. SOX18^+^/SREBP2^+^ double positive cell nuclei in involuting phase IH specimens were significantly reduced compared to proliferating phase IH (Figure 4A,B,C, Supplemental Figure 4.1-3). Moreover, we analyzed tissue from two patients with regrowing IH. Both patients received beta-blocker treatment during infancy with notable clinical response. However, the IH regrew significantly after discontinuation of treatment. Both regrowing IH specimens were positive for nuclear SOX18 and SREBP2 along the endothelium. We observed histological features of proliferating IH (nuclear co-localization of SOX18 and SREBP2) as well as involuting IH (enlarged vessels, connective tissue with stromal cells and adipocyte islands) in the regrowing IH (Figure 4D, Supplemental Figure 4). Anti-SOX18 staining specificity was verified as shown (Supplemental Figure 4.5-8). Quantification of SOX18^+^SREBP2^+^ cells/total endothelial cell nuclei in regrowing IH demonstrated a similar level as seen in proliferating IH and significantly increased compared to involuting IH and age-matched skin controls (Figure 4E). In summary, the nuclear co-localization of SOX18 and SREBP2 in proliferating and regrowing phase IH underscores the role of the MVP in IH pathogenesis and may serve as markers to predict therapy response.

**Figure 4.**
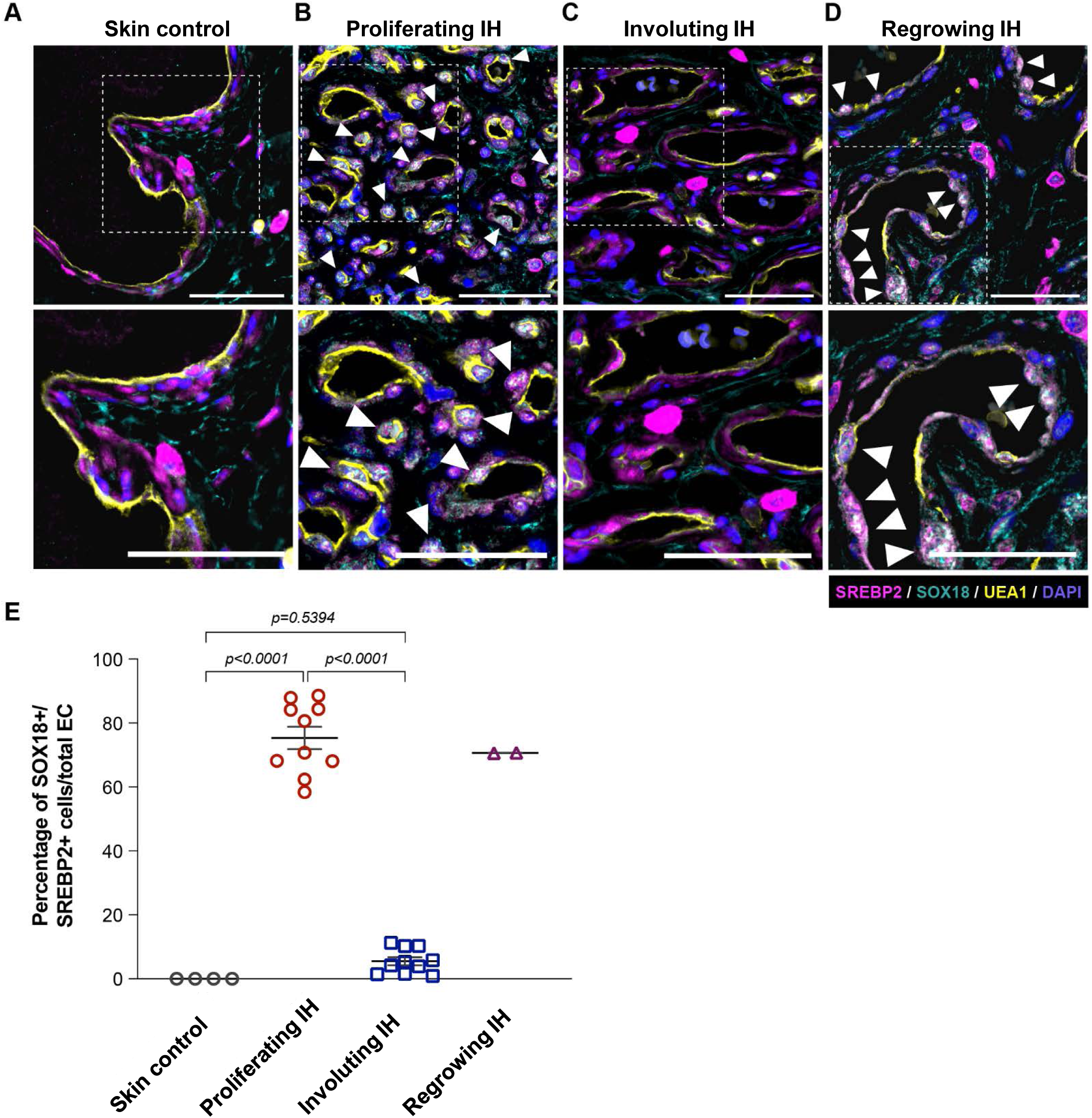
Nuclear co-localization of SOX18 and SREBP2 in proliferating phase and regrowing IH indicate active MVP. (**A**-**D**) Human age-matched skin, proliferating IH, involuting IH and regrowing IH stained for SREBP2 (magenta), SOX18 (cyan), and the human EC-specific lectin UEA1 (yellow). Cell nuclei are stained with DAPI (blue). SOX18^+^/SREBP2^+^ double positive cell nuclei (arrowheads) are significantly more abundant in proliferating phase and appear increased in regrowing compared to involuting phase IH and age-matched skin controls, as quantified in (**E**). P values were calculated using one-way ANOVA with Šidák-correction. Data show the mean ± SD. Data represent sample sizes for n=4 age-matched skin controls, n= 10 each for proliferating and involuting phase IH, and n=2 for regrowing IH. Scale bars 50 μm.

### Statins inhibit vessel formation in a xenograft model of IH

We surmised that if the MVP is critical to IH onset and progression, statins would inhibit HemSC blood vessel formation in an IH preclinical model. The effect of statins on patient-derived cells has not been reported before and we therefore first tested for cell toxicity and efficacy of statins on HemSC. We chose to use simvastatin and atorvastatin for our preclinical model as they are commonly used statins in patients, even for infants in the case of simvastatin (31). It is well established that atorvastatin is 5-10 times more potent than simvastatin (32). To first address toxicity, HemSC (n=3) were induced to undergo endothelial differentiation until day 6 while treated with simvastatin (0.1 - 1 μM) or atorvastatin (0.01 - 0.1 μM) and respective vehicle controls. Neither statin had an effect on differentiating HemSC viability compared to vehicle controls (Supplemental Figure 5.1). Next, we tested the effect of statins on LDL-receptor levels in HemSC as an indirect measure of their effect on the MVP. Treatment with 5 μM simvastatin or 1 μM atorvastatin for 24 hours resulted in a significant upregulation of LDL-R mRNA in HemSC (n=4) (Supplemental Figure 5.2), indirectly indicating efficient inhibition of HMGCR in these cells.

We first tested 1 μM simvastatin and 0.1 μM atorvastatin on VEGF-B-induced HemSC endothelial differentiation as described(8). Both simvastatin and atorvastatin significantly inhibited endothelial differentiation as indicated by decreased expression of the EC markers CD31 and VE-Cadherin compared to vehicle control (n=3). R(+) propranolol was included as a positive control (Figure 5A). We next tested if statins would impact de novo vessel formation in the murine xenograft model using IH-derived HemSC (n=4 different patients) (Figure 5B). Simvastatin at 10 mg/kg/day (d) (Figure 5C,D) and atorvastatin at 1 mg/kg/d (Supplemental Figure 5.3) both significantly inhibited human CD31+ blood vessel formation. A simvastatin dose response experiment further showed that 1 mg/kg/d was sufficient to significantly inhibit vessel formation (Figure 5E). Glucose levels and body weight of mice in the simvastatin dose response experiment were unaffected (Figure 5F,G). The human equivalent doses corresponding to the doses used in mice are shown in Supplemental Figure 5, Supplemental Table 5.1. The effective doses of simvastatin and atorvastatin in Figure 5 are below effective doses used in adults and significantly below the dose used in infants for Smith Lemli Opitz Syndrome (0.5 - 1 mg/kg/d simvastatin). Of note, the inhibitory effect of statins was limited to HemSC de novo vessel formation and did not impact angiogenic sprouting and ingrowth of surrounding murine vessels into the Matrigel implant (Supplemental Figure 5.4-5). Specificity of the anti-human CD31 and anti-mouse CD31 antibodies was verified (Supplemental Figure 5.6). Taken together, our data show that simvastatin and atorvastatin inhibit endothelial differentiation of HemSC in vitro and vasculogenesis in a preclinical IH xenograft model. This strongly suggests the MVP contributes to vascular overgrowth in IH and can be effectively targeted by statins in a translational approach.

**Figure 5.**
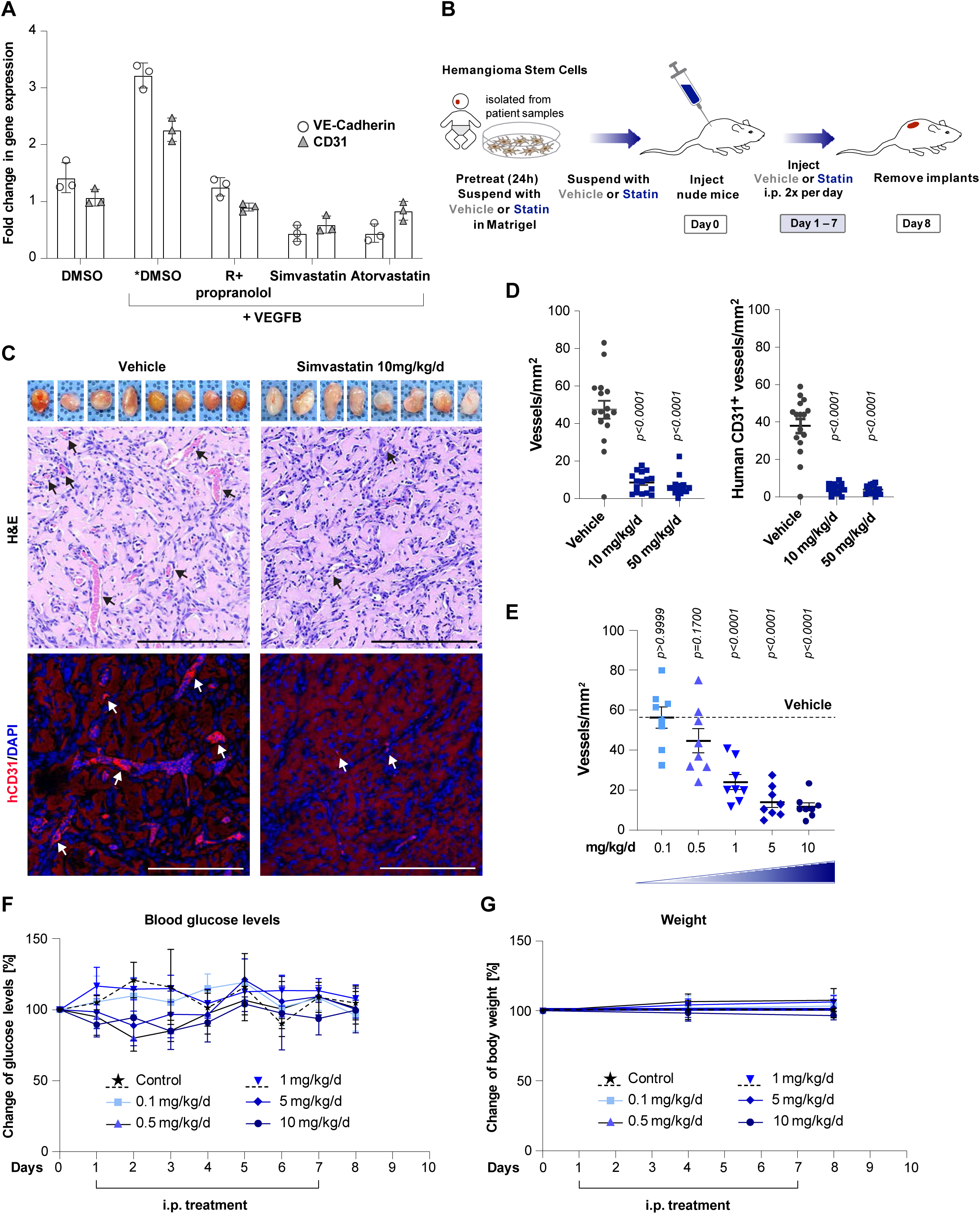
Statins inhibits HemSC endothelial differentiation and blood vessel formation in a xenograft model of IH. (**A**) Simvastatin (1 μM) and atorvastatin (0.1 μM) inhibited VEGF-B induced endothelial differentiation of HemSC (n=3) in vitro as indicated by reduced mRNA levels of EC markers VE-Cadherin and CD31 over the course of 6 days of treatment. R(+) propranolol (20 μM) served as a positive control. P values shown in Supplemental Table S5.2) (**B**) Schematic of IH xenograft model. Patient-derived HemSC (n=4) were pre-treated with 5 μM simvastatin or vehicle (0.005% DMSO), suspended in Matrigel with 2.5 μM simvastatin or vehicle (0.0025% DMSO), and injected subcutaneously into nude mice, with 2 implants/mouse. Mice were treated with 0.1 - 50 mg/kg/d simvastatin and vehicle (max. 2.3% DMSO/dose) twice a day via intraperitoneal injections. (**C**) Matrigel implants harvested after 7 days of treatment are shown in the top panel. H&E staining indicates reduction of blood vessels in simvastatin treated compared to control mice (middle panels). Anti-human CD31 staining (red) confirmed reduced human vessel formation in simvastatin-treated mice compared to controls (bottom panels). Cell nuclei were stained with DAPI (blue). Scale bars 100 μm. (**D**) Quantification of vessels/mm^2^ in the H&E-stained sections (left) and human CD31^+^ vessels/mm^2^ (right) show a significant reduction in vessel density in the implants of simvastatin-treated mice compared to control mice. (**E**) Dose response to simvastatin indicates significant inhibition of vessel formation as of 1 mg/kg/d. (**F**,**G**) Blood glucose levels and body weight of mice were unaffected in both simvastatin-treated and control mice. P values were calculated using one-way ANOVA multiple comparisons test with Tukey-correction (**A**,**D**), one-way ANOVA multiple comparisons test with Dunnett-correction (**E**), and two-way ANOVA multiple comparisons test with Dunnett-correction (**F**). Data are shown as ± SD. Data were collected for 2 implants in each mouse, leading to an observation sample size of n=24 for vehicle (combined), n=14 for 10 mg/kg/d, and n=16 for 50 mg/kg/d simvastatin (**D**) as well as in the dose response n=10 for vehicle, n=10 for 10 mg/kg/d, n=10 for 5 mg/kg/, n=10 for 1 mg/kg/d, n=10 for 0.5 mg/kg/d, and n=10 for 0.1 mg/kg/d simvastatin (**E**).

## DISCUSSION

In this study, we uncovered the SOX18-MVP axis as a central regulator of IH pathogenesis. The R(+) enantiomer of propranolol, shown previously to disrupt SOX18 transcriptional activities, downregulates expression of several genes in the MVP in a SOX18 dependent manner and reduces mature SREBP2 protein, the master regulator of the MVP. R(+) propranolol or the SOX18 inhibitor Sm4 decreased new synthesis of cholesterol in HUVEC. We propose that SOX18 augments the transcription of MVP genes to regulate cholesterol and isoprenoid biosynthesis in HemSC undergoing endothelial differentiation. Of note, SOX18 levels increase over the course of HemSC to EC differentiation; this corresponds to MVP sensitivity to R(+) propranolol. Furthermore, *SOX18* ChIP binding sites are present in *HMGCS1* and *HMGCR*. Together, our experimental results and observations make a compelling case for the involvement of the SOX18-MVP axis in the etiology of IH and suggests statins may be a new therapeutic option.

Additional experiments support these novel insights. Competitive inhibition of the rate-limiting enzyme of the MVP - HGMCR - with statins (simvastatin and atorvastatin) results in significantly reduced HemSC blood vessel formation in mice in a dose-dependent manner with unaffected weight and glucose levels. Furthermore, the numbers of angiogenic mouse blood vessels in the HemSC/Matrigel implants were unaffected, indicating statins have little or no effect on host angiogenesis. Additional evidence for SOX18 regulation of the MVP was obtained from *RaOp* SOX18 expressing HUVEC. The *RaOp* dominant negative SOX18, which is thought to “poison” normal SOX18 function (33), downregulated HMGCS1 and HMGCR. SOX18 and SREBP2 were found to colocalize in endothelial nuclei in proliferating phase IH and two cases of regrowing IH but not in involuting phase IH or normal skin indicating nuclear SOX18 and SREBP2 may be restricted to nascent vessels in IH. Regrowing IH defies the classic IH life cycle, and little is known about how or why these IH regrow; most often, regrowing IH involves oral mucosa, such as the lip. Typically, propranolol therapy is restarted to prevent sequelae. In our two IH patients, regrowth occurred at 6 and 9 years of age. The nuclear co-localized SOX18 and SREBP2 in proliferating phase IH and regrowing IH suggests that SOX18^+^/SREBP2^+^ endothelial nuclei may serve as a biomarker for the vasculogenic capacity of the tumor. This may give important biomolecular clues for patients to predict treatment response to statins and may thereby provide a novel theranostic approach.

SOX18 is a firmly established transcriptional regulator of vascular and lymphatic development, and tumor angiogenesis (11–14, 25, 34). The small molecule inhibitor of SOX18, Sm4, suppresses vascular development in zebrafish and both tumor angio- and lymph-angiogenesis in a breast cancer model (25, 26). Notably, while glycolysis and fatty acid oxidation have been well established in physiological and pathological endothelial cell metabolism (35, 36), the MVP has not been implicated as a crucial regulatory pathway. In line with R(+) propranolol downregulation of SQLE (Figure 1C, Supplemental Table 1), the gene encoding squalene epoxidase, a key enzyme in cholesterol biosynthesis, single cell RNA sequencing of endothelial cells in pathological angiogenesis identified SQLE as a metabolic angiogenic target (37). Our discovery of a functional link between SOX18 and the MVP brings to light an entirely novel concept in endothelial differentiation and vasculogenesis. By interfering with SOX18 transcriptional activity and downstream regulation of the MVP, R+ propranolol acts as an EC phenotype-specific “statin”.

The MVP is central to cholesterol biosynthesis and isoprenoid biosynthesis, the later needed for prenylation. The MVP bifurcates at farnesyl pyrophosphate (FPP) to produce squalene, an intermediate in cholesterol biosynthesis, or geranylgeranyl pyrophosphate (GGPP). FPP and GGPP are directly involved in prenylation of various small GTPases, and other protein substrates including RAS family members (38). Importantly, RAS signaling has been implicated in IH (39). The R(+) propranolol mediated reduction in MVP genes and cholesterol biosynthesis may affect membrane fluidity and ruffling or activation of important signaling pathways by prenylation. This warrants further investigation.

Pleiotropic benefits of statins, beyond their lowering effect on plasma LDL-cholesterol levels, are well documented and thus inferred to provide cardiovascular disease protection. These include inhibiting inflammatory response, increasing the bioavailability of nitric oxide, promoting re-reendothelialization, and reducing oxidative stress (19, 20, 40). The underlying mechanisms, however, remain elusive. A recent study suggests epigenetic effects: simvastatin significantly improved human induced pluripotent stem cell-derived endothelial cell function by reducing chromatin accessibility under physiological and pathological conditions (24). Moreover, statins block mammalian target of rapamycin (mTOR) and thereby inhibit AKT signaling (41), which is central to balancing vascular growth with endothelial quiescence. It was further shown that mTOR complex 1 reduces ER cholesterol levels which in turn activates SREBP2 and the MVP (42). Of note, PI3K-AKT-mTOR signaling is crucial in the pathogenesis of slow-flow vascular malformations (43, 44). Hence, statins may hold promise to be efficient in these vascular anomalies as well and may be relevant for future directions.

Statins are widely used drugs. The resulting reduction in cholesterol synthesis consequently increases LDL-receptor expression, which in turn, clears low-density lipoprotein cholesterol from the circulation. Although statins are generally considered a safe and well tolerated drug class, they are associated with an increased risk of muscle pain, diabetes mellitus and hepatic transaminase elevations (45). We addressed potential side effects by measuring glucose levels and body weight in the statin treated xenograft mice. We did not observe a change in glucose levels or weight with simvastatin and atorvastatin over the treatment course of 7 days. This is a relatively short treatment duration to detect adverse effects and moreover, side effects in rodents differ from those in humans. Bearing in mind drug safety as the highest priority when considering translating statins to infants, we performed a dose response experiment with simvastatin. The lowest dose that showed significant reduction in IH vessels in the xenograft model was 1 mg/kg/d (Figure 5). This translates to a human equivalent dose of 0.081 mg/kg/d (46) (Supplemental Figure 5, Table S5.2), which is 6.25 times below the recommended dose of 0.5 mg/kg/d systemic simvastatin for infants with other indications. Further, topical rather than systemic statins might be used for superficial or less complex IH with increased safety as shown for other applications discussed below.

Systemic statins have been used safely in infants with Smith-Lemli-Opitz syndrome (SLOS). In a randomized, placebo-controlled clinical trial the recommended dose was 0.5 to a maximum of 1 mg/kg/d. SLOS is an autosomal-recessive multiple malformation/cognitive impairment syndrome and characterized by the accumulation of 7-dehydrocholesterol, a precursor of cholesterol (31, 47–49).

Topical statins have also been used for dermatological conditions such as congenital hemidysplasia with ichthyosiform erythroderma and limb defects (CHILD) syndrome - an X-linked, dominant condition with most surviving patients being female (50). In addition, statins have been used to treat alopecia areata (51, 52). Based on the successful use of statins in children with these disorders, we suggest a safe repurposing of topical and systemic simvastatin to treat infants with IH.

It is exciting to speculate that the novel link between SOX18 and the MVP may have important implications for various pathophysiological conditions of the endothelium, including developmental defects in vascular anomalies and other disease scenarios with disturbed vasculogenesis and angiogenesis. In contrast to vascular malformations for which genetic drivers have been identified in recent years, the genetic basis of IH is unknown. The advanced understanding of genetics has facilitated targeted therapies for vascular malformations translating pre-existing oncology drugs that interfere with signaling pathways involved (53, 54). Conversely, critical molecular players in IH have been uncovered by studying mechanisms of action of serendipitously discovered drugs (8, 9, 55, 56). This has led to our discovery that demonstrates a mechanistic link between beta-blockers and statins via the endothelial cell specific transcription factor SOX18.

We propose the SOX18-MVP-axis may connect broadly to clinical studies that link beta-blocker and statin use to improved outcomes in cancer patients. We speculate this is due to suppression of the SOX18-MVP-axis in tumor endothelial cells and in turn, tumor angiogenesis. Propranolol is currently being studied as single administration adjuvant therapy or in combination with standard therapies in several clinical trials (57). Statin use was first found associated with improved outcomes in colorectal cancer patients in 2005 (58) and subsequent studies confirmed this observation and extend to post-diagnosis statin use (59–64).

In summary, our findings uncover a novel SOX18-MVP axis as a central molecular driver for HemSC vasculogenesis. In a xenograft model we show that simvastatin or atorvastatin inhibit IH vessel formation. This suggests statins could be repurposed to treat this common vascular tumor of infancy, topically or systemically. Moreover, we demonstrate suppression of the transcription of MVP genes via R+ propranolol disruption of SOX18. This provides a mechanistic link between two of the most widely used drugs in cardiovascular disease – beta-blockers and statins. We propose that the SOX18-MVP axis may explain the pleiotropic effects of beta-blockers and statins in a wide range of clinical scenarios that involve pathological vasculogenesis and angiogenesis.

## METHODS

### Sex as a biological variable

Our study used male mice for xenograft of hemangioma-derived cells isolated IH from male and female patients (see Table 1).

### IH cell isolation and culture

The clinical diagnosis of IH was confirmed in the Department of Pathology of Boston Children’s Hospital. Single-cell suspensions prepared from proliferating IH specimens were de-identified and designated as specified in Table 1. HemSC or HemEC were selected using anti-CD133– and anti-CD31–coated magnetic beads (Miltenyi Biotec), respectively, and expanded. Testing for mycoplasma contamination by qPCR was performed when cells were thawed and every 4–6 weeks thereafter. Cells were cultured on fibronectin-coated (0.1 μg /cm^2^; MilliporeSigma) plates with a density of 20,000 cells/cm^2^ with Endothelial Growth Medium-2 (EGM-2; Lonza), which consists of Endothelial Growth Basal Medium-2 (EBM-2; Lonza), SingleQuots supplements (all except hydrocortisone and gentamicin-1000; Lonza), 10% heat-inactivated FBS (Cytiva), and 1× GPS (292 mg/mL glutamine, 100 U/mL penicillin, 100 mg/mL streptomycin, Mediatech). Hereafter, this full growth media is referred to as EGM-2. Cells were cultured at 37°C in a humidified incubator with 5% CO2 and fed every other day.

### Hemangioma endothelial differentiation assay

To induce endothelial differentiation, HemSC (samples denoted as HemSC 125, 149, 150, 165, 167, 171) were seeded on fibronectin-coated plates (0.01 μg fibronectin/cm^2^) at a density of 20,000 cells/cm^2^ in EGM-2. After 18 - 24 hours, cells were starved for 16 hours in 2% BSA/serum-free EBM-2. Cells were washed with EBM-2 once. The medium was replaced by differentiation media that consisted of serum-free EBM-2 containing 1× insulin transferrin-selenium, 1:100 linoleic acid–albumin, 1 μM dexamethasone, and 100 μM ascorbic acid-2-phosphate. The medium was replaced with fresh differentiation medium containing 10 ng/mL VEGF-B (R&D Systems) every two days. For addition of inhibitors, a preincubation ± respective inhibitors (R(+) propranolol, simvastatin, or atorvastatin) for 30 minutes was conducted followed by continuous treatment. Stock solutions of 10 mM R(+) propranolol hydrochloride (MilliporeSigma), 105 mM simvastatin, and 10mM atorvastatin (both Selleckchem) were prepared in DMSO (MilliporeSigma). 20 μM R(+) propranolol was added for two hours and cells harvested (n=6) for RNA sequencing (Figure 1A,B,C). Continuous (4 consecutive days) as well as short term treatment (2 hours) with 20 μM R(+) propranolol was applied to validate the RNA sequencing data of HemSC undergoing endothelial differentiation by qPCR (n=3) (Figure 1D,E).

Continuous treatment with 1 μM simvastatin, 0.1 μM atorvastatin as well as 20 μM R(+) propranolol as a positive control was applied to differentiating HemSC (n=3) to test for an effect of statins in vitro (Figure 5A). Respective vehicle controls of each drug were used. Vehicle without VEGF-B served as a negative control and with VEGF-B as a positive control for differentiation. A potential toxic effect of simvastatin and atorvastatin on viability of HemSC (n=3) ± simvastatin, atorvastatin, or vehicle incubated for 24 hours was assessed by cell counting (Supplemental Figure 5.1).

### Cholesterol measurements in HUVEC

HUVEC were grown in EBM-2 media (Lonza, CC-3156) and supplemented with EGM-2 bullet kit (Lonza, CC-3162). HUVEC were seeded on 0.5% gelatin-coated 6-well plates at a density of 1.5 x10^5^ cells/well overnight. The day after seeding cells were treated with MBCD (Sigma C4555) at a concentration of 2.5 mM to deplete cells of endogenous cholesterol. MBCD was withdrawn after 4 hours and cells were washed with PBS. After this, cells were treated with either PBS, 20 μM R(+) propranolol (Sigma-Aldrich, Cat P0689), DMSO, or 40 μM of Sm4 (Sigma-Aldrich, SML1999) for 18 hours. Cells were then washed with PBS, trypsinized, washed with PBS, and centrifuged (Figure 2B,C). Lipids were extracted from the cell pellets through one-phase extraction with 202 uL of BuOH:MeOH (1:1) (Sigma 71-36-3 and 67-56-1) that contains 10 umol of cholesterol-d7 (Cayman Chemical 83199-47-7) as the internal standard (65). The cholesterol analysis was performed on a TSQ Altis triple quadrupole mass spectrometer, operated in positive ion mode, coupled to a Vanquish UHPLC system (Thermo Fisher, Waltham, USA) using the transition from precursor mass of *m/z* 369.3516 (MS1) to *m/z* 161.1 (MS3) for cholesterol and 376.4 to 161.1 for cholesterol-d7 (65). The solvent pair included solvent A (100% H2O, 0.1% FA and 2 mM NH4COO^-^) and solvent B (100% MeOH, 0.1% FA and 2 mM NH4COO^-^) with a flow rate at 0.3 mL/min (80% B at the start). Cholesterol and its deuterated standard were separated on an Eclipse Plus C8 column (Agilent), and peaks were integrated with TraceFinder 5.1 (Thermo Fisher, Waltham, USA) using the daughter ion at *m/z* 161.1 (Figure 2B,C) (66).

### RaOp expression in HUVECs

The day after HUVEC seeding, transfection was performed using X-tremeGene HP Transfection Reagent kit (Roche) to introduce 500ng of Halo-RaOp DNA using EBM-2 culture media without antibiotic. Cells were incubated at 37°C with 5% CO2 overnight. The next day, cells were incubated with 5nM of JF646-Halo-ligand for 15min, washed with PBS and trypsinized; as a result, HUVEC expressing SOX18*^RaOp^* were fluorescently tagged. Cells were spun down, washed with PBS and permeabilized with 0.2% Triton X-100 for 10min, following blocking with 5% BSA/PBS for 1 hour. Cells were labelled with 1:300 dilution of anti-human HMGCR (ThermoFisher, PA537367) or HMGCS1 (ThermoFisher, PA513604) for 1 hour at room temperature in the dark. Cells were washed with PBS and stained with 1:500 dilution of anti-rabbit IgG conjugated with Alexa Fluor 488 (Invitrogen A11008) for 30 min and washed with PBS. Cells were then imaged on a Fortessa X-20 (BD Bioscience). Data analysis was performed using FlowJo software (Figure 2D,E).

### shRNA knockdown of SOX18

HemEC (n=3) were transduced with SOX18 shRNA lentivirus (TRCN0000017450, Sigma) or an empty vector control virus (Sigma SHC001V) followed by puromycin (1 μg/mL) selection for 5 days. Thereafter, cells were maintained in EGM-2 without puromycin. Knockdown efficiency was confirmed by qPCR and Western Blot (Figure 3C-G, Supplemental Figure 3.1,2).

### Western Blot

HemSC were seeded on fibronectin-coated plates (0.1 μg fibronectin/cm^2^) at a density of 20,000 cells/cm^2^ in EGM-2. HemSC were induced to undergo differentiation according to the protocol described above. Cells were treated with 20 μM R(+) propranolol for 2 hours on Day 6 of endothelial differentiation and lysed in a RIPA-based lysis buffer (25 mM Tris HCl pH 7.6, 150 mM NaCl, 1% NP-40, 1% sodium deoxycholate, 0.1% SDS) with protease and phosphatase inhibitors (Cell Signaling Technology, 5872S) as well as a calpain I and proteasome V inhibitors (Millipore Sigma, 208719). Cell extracts were electrophoresed by sodium dodecyl sulfate-polyacrylamide gel electrophoresis (SDS-PAGE), transferred to nitrocellulose or to polyvinylidene difluoride (PDVF) and probed with anti-SREBP2 clone 22D5 (Millipore Sigma, MAB1988) followed by anti-GAPDH (Cell Signaling Technology, 5174).

For protein quantification of shSOX18 knockdown efficiency in HemEC^shSOX18^ (samples denoted as HemEC ^shSOX18^ 133, 150 and 171), cells were seeded on fibronectin-coated plates (0.1 μg fibronectin/cm^2^) at a density of 20,000 cells/cm^2^ in EGM-2 media, lysed and processed as described above following staining with SOX18 D-8 (Santa Cruz Biotechnology, sc-166025) and GAPDH (Cell Signaling Technology, 5174) antibodies. All signals were detected by enhanced chemiluminescence. The densitometric analysis was conducted using Fiji ImageJ software.

### In vivo murine model for human blood vessel formation

In vivo, HemSC undergo vasculogenesis and form anastomoses with ingrowing host vessels. Experiments were carried out with 3 × 10^6^ HemSC per implant. HemSC (n=4, samples 125, 147, 149, 150) were grown in EGM-2 media until 90% confluency. A stock solution of simvastatin (105 mM; MilliporeSigma) or atorvastatin (10 mM; MilliporeSigma) was prepared in DMSO. The stock solution was diluted with PBS to the indicated treatment concentrations. Twenty-four hours before harvesting, 5 μM treatment with simvastatin or 0.5 μM treatment with atorvastatin or the equivalent DMSO concentration (0.005%) as a control was added to the media. Cells were counted after the 24-hour pretreatment and suspended in 200 μL Matrigel (Corning) preadjusted with 1 μg/mL basic FGF (bFGF) (ProSpec), 1 μg/mL erythropoietin (EPO) (ProSpec), and 2.5 μM (simvastatin) or 0.25 μM (atorvastatin) drug or PBS with max 0.005% DMSO on ice. The Matrigel/cell suspensions were injected subcutaneously into the flanks of 6-week-old male athymic nu/nu mice (envigo), placing 2 implants per mouse (n = 3-5 mice/group; Figure 5B). The mice were treated with 0.1 - 50 mg/kg/d simvastatin, 1 - 15 mg/kd/d atorvastatin or the equivalent DMSO concentration (maximal 11.4 %) as a control (200 μL/mouse, i.p.) twice a day. Blood glucose levels of the mice were measured daily before the morning i.p. injection. Glucose concentrations were measured in tail vein blood using the OneTouch UltraSmart Blood Glucose Monitoring System (LifeScan). Body weight was measured before the injections (day 0), on day 4 and before removal of the implants (day 8). After 8 days, the mice were euthanized and the implants were removed, fixed in formalin, embedded in paraffin, and analyzed by H&E staining and immunofluorescent staining (IF). Blood vessels (indicated by luminal structures containing one or more red blood cells) and CD31^+^ stained human vessels were counted in 5 fields/section, 2 sections/implant. Each field was 425.1 μm × 425.1 μm = 0.18071 mm2, and sections were from the middle of the implant. Vessel density is expressed as vessels/mm^2^.

### Histology and IF analysis

FFPE tissue sections (5 μm) of the Matrigel implants were deparaffinized and either directly stained with H&E or immersed in an antigen retrieval solution (citrate-EDTA buffer containing 10 mM citric acid, 2 mM EDTA, and 0.05% Tween-20, pH 6.2) for 20 minutes at 95°C–99°C. Sections were subsequently blocked for 30 minutes in TNB Blocking Buffer (PerkinElmer) followed by incubation with an anti-human CD31 monoclonal antibody (1:30, mouse anti-human; Dako, Glostrup, 0823) to stain for human endothelium. Next, sections were incubated with Alexa Fluor 647 chicken anti-mouse IgG (1:200, Invitrogen, Thermo Fisher Scientific) as a secondary antibody (Figure 5C,D).

An anti-mouse-specific CD31 monoclonal antibody (1:100, R&D Systems,) was used to quantify mouse vessels in the Matrigel implants. Alexa Fluor 647 chicken anti-goat IgG was applied as a secondary antibody (Supplemental Figure 5.4). Tissue specificity of the anti-human and anti-mouse antibodies was confirmed by negative staining in mouse lung and human skin tissue, respectively (Supplemental Figure 5.6).

FFPE tissue sections (5 μm) from patients with IH were deparaffinized, immersed in an antigen retrieval solution, and blocked for 30 minutes in 10% donkey serum followed by incubation with mouse anti-human SOX18 (1:50; Santa Cruz Biotechnology), rabbit anti-human SREBP2 and UEA1 fluorescently labeled with Alexa Fluor 649 (1:50; Vector Laboratories). Next, the sections were incubated with Alexa Fluor 488 donkey anti-mouse IgG and Alexa Fluor 546 donkey anti-rabbit IgG (both 1:200; Invitrogen, Thermo Fisher Scientific) as secondary antibodies. All slides were mounted using DAPI (Molecular

Probes, R37606) to visualize nuclei. IF Images were acquired by a LSM 880 confocal microscope (Zeiss). Images were analyzed through a 20x or 63x objective lens. All images were analyzed using Fiji ImageJ software (Figure 4).

### Cell staining

HemEC171^naïve^ and HemEC171^shSOX18^ were seeded on fibronectin-coated plates at a density of 20,000 cells/cm^2^ in EGM-2 media for 30 hours on 2 cm^2^ slides, fixed in 4% PFA and blocked in 5% BSA/0.3% Triton x-100 for 1 hour. The mouse anti-human SOX18 primary antibody used for immunostaining (E-11) was validated in HemEC^naïve^ and HemEC^shSOX18^ (1:100, Santa Cruz, sc-376166), Cells were co-stained with a rabbit anti-human VE-Cadherin (1:100, R&D Systems). Secondary antibodies included Alexa Fluor 488 donkey anti-mouse IgG and Alexa Fluor 546 donkey anti-rabbit IgG (both 1:200; Invitrogen, Thermo Fisher Scientific). Slides were mounted using DAPI (Molecular Probes, R37606) to visualize nuclei (Supplemental Figure 4.5,6). Immunostaining of IH sections with respective isotype matched control IgG and secondary antibodies are shown in (Supplemental Figure 4.7,8).

### RNA isolation and qPCR

Total cellular RNA was extracted from cells with a RNeasy Micro Extraction Kit (QIAGEN). Reverse transcriptase reactions were performed using an iScript cDNA Synthesis Kit (Bio-Rad). qPCR was performed using SYBR FAST ABI Prism 2× qPCR Master Mix (Kapa BioSystems). Amplification was carried out in a QuantStudio 6 Flex Real-Time PCR System (Thermo Fisher Scientific). A relative standard curve for each gene amplification was generated to determine the amplification efficiency, with greater than 90% considered acceptable. Fold increases in gene expression were calculated according to the ΔΔCt method, with each amplification reaction performed in duplicate or triplicate (67). Gene expression was normalized to the PBS treatment. ATP5B was used as housekeeping gene expression reference.

### Bioinformatic analysis

#### Bulk RNA-seq

We used trimmomatic v0.39 (68) to trim the low-quality next generation sequencing (NGS) reads (-threads 20 ILLUMINACLIP:TruSeq3-PE.fa:2:30:10 LEADING:3 TRAILING:3 SLIDINGWINDOW:4:15 MINLEN:36). Subsequently, only the high-quality trimmed reads were aligned to the human reference genome (hg38) using STAR v2.7.2b. The reads counts were calculated by featureCounts software (69). Differentially expressed genes (DEGs) were identified by using the DESeq2 R package (adjusted p-value < 0.05). KEGG pathways and gene ontology (GO) enrichment tests were performed by the clusterProfiler R package. A pathway or GO term was treated as significantly enriched if an adjusted p-values (with Benjamini-Hochberg correction) was smaller than 0.05. The bar plots illustrating significant pathway or GO terms were created using the enrichplot R package.

#### ChIP-seq

The data is displayed on the UCSC genome browser. The layered epigenetic marks dataset is supplied from Encode. ChIP-seq binding location dataset can be found at https://www.ebi.ac.uk/biostudies/arrayexpress/studies/E-MTAB-4481.

#### Statistics

Data were analyzed and plotted using GraphPad Prism 10.1.0 (GraphPad Software). Results are displayed as the mean ± SD. For experiments in which cells were treated with alternative drugs, the differences were assessed by one-way ANOVA. Tukey’s post hoc test was used for multiple comparisons of different treatment modalities and Šidák’s, Tukey’s or Dunnett’s test for multiple comparisons to compare every treatment mean with that of the vehicle control. A two-tailed, unpaired t-test was applied for comparisons between treatment and control groups given equal variance.

#### Study approval

Animal protocols complied with NIH Animal Research Advisory Committee guidelines and were approved by the Boston Children’s Hospital Animal Care and Use Committee (protocol number 00001741). IH specimens were obtained under a protocol approved by the Committee on Clinical Investigation at Boston Children’s Hospital (IRB protocol number 04-12-175R; AH, HK, JBM, JB) as well as at Stanford University (IRB protocol number 35473; RL, JT). Hemangioma specimens were collected upon written informed consent of the patient’s guardian, deidentified, and used for cell isolation under a Boston Children’s Hospital IRB approved protocol (04-12-175R) and in accordance with Declaration of Helsinki principles.

## Acknowledgements

We thank the Vascular Anomalies Center and Department of Pathology at Boston Children’s Hospital for clinical care and for sharing patient specimens.

We thank Trisha Lopez for patient chart reviews, Dr. Sandra Schrenk for her intellectual input and technical advice, Dr. Andrew Stone for imaging advice and Kristin Johnson of the Vascular Biology Program graphics core for figure design. Research reported in this manuscript was funded by NIH grant 5R01HL096384-012 (to JB). MF, MG are supported by the National Health and Medical Research Council, GNT1164000. AH was supported by the Deutsche Forschungsgemeinschaft (DFG, German Research Foundation, grant 458322953), and is supported by the Vascular Anomalies Center at Boston Children’s Hospital. JWHT received funding from the Nanyang Technological University CN Yang Scholars Program, AH and MV are members of the Vascular Anomalies Working Group (VASCA) of the European Reference Network for Rare Multisystemic Vascular Diseases (VASCERN) - Project ID: 769036.

The funders had no role in the study design, data collection and interpretation, or decision to submit this work for publication.

## Author contributions

AH, MG, MF and JB designed the study. AH, MG, JWS, LB, LC, AJ, MS, LS, MF and JB designed the methodology. AH, MG, JWS, LB, JWHT and LC performed functional investigations. AH, MG, JWS, LB, JWHT, LC, AJ, MS, and LS carried out formal analysis. AH and JB wrote the original draft. All authors performed reviewing and editing. All authors read and agreed to the final version of the manuscript. AH performed patient data curation. RL, HK, JCT and JBM provided patient specimens and clinical expertise. LS, JQ, JCT, MV, MF and JB provided supervision. MF and JB conceived the study. AH, JWHT, MV, MF and JB acquired funding.

## Intellectual Property

AH and JB are co-inventors on a filed patent application PCT/US23/83306 “Methods and Compositions for the Treatment of Vascular Anomalies.” JB is co-inventor on

US Patent # 9,737,514.

