## Supplemental Figures for "An endothelial SOX18-mevalonate pathway axis enables repurposing of statins for infantile hemangioma"

### Supplemental Figure 1

## S1.1

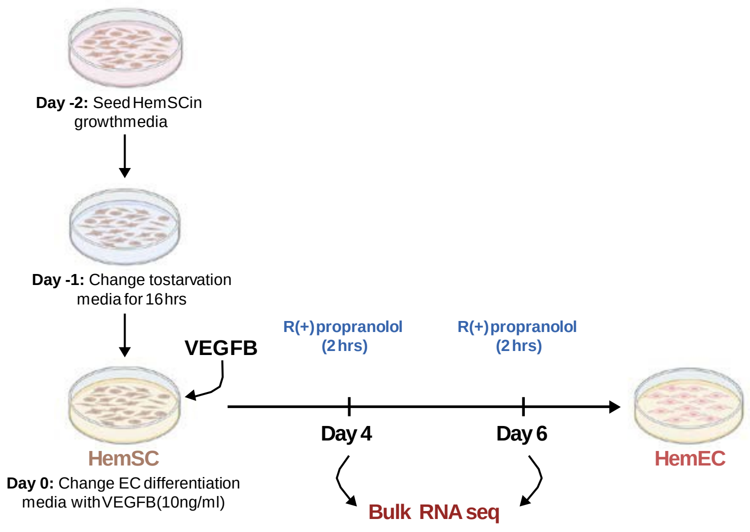


## S1.2

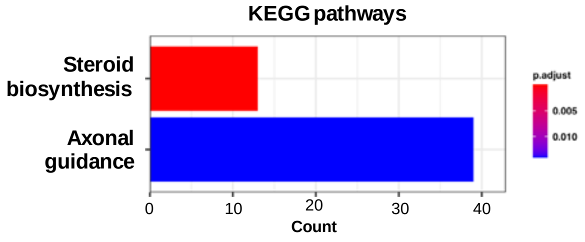


#### Table S1

| | Day 4 | | Day 6 | |
|---|---|---|---|---|
| **Gene ID** | **Log$_2$ fold change** | **Adjusted p value** | **Log$_2$ fold change** | **Adjusted p value** |
| ***HMGCS1*** | 0.462979006 | 0.026354098 | -1.63371667 | 0.000484519 |
| ***HMGCR*** | 0.169344377 | 0.044323597 | -1.374323007 | 0.000424613 |
| ***MVK*** | 0.314895512 | 0.029933298 | -1.147210743 | 0.000127633 |
| ***MVD*** | 0.441665738 | 1.75E-06 | -1.288125115 | 0.002855957 |
| ***FDPS*** | 0.217777666 | 0.019564343 | -1.033766375 | 0.006204092 |
| ***IDI1*** | 0.24599104 | 0.010011861 | -1.226584676 | 0.004495371 |
| ***FDFT1*** | 0.173687183 | 0.080502823 | -0.874992512 | 0.011617406 |
| ***SQLE*** | 0.186643474 | 0.080502823 | -1.274290854 | 0.001552676 |
| ***LSS*** | 0.161258298 | 0.088436956 | -0.74862184 | 0.017189697 |
| ***SC5D*** | 0.079030565 | 0.728721351 | -0.96705494 | 0.005332315 |
| ***HSD17B7*** | 0.211229037 | 0.550897945 | -1.81198247 | 2.01E-06 |
| ***NSDHL*** | 0.192680874 | 0.220906602 | -0.686486501 | 0.015844376 |
| ***DHCR7*** | 0.231028296 | 0.09552771 | -1.404321837 | 0.002936361 |
| ***DHCR24*** | 0.039092468 | 0.885153473 | -1.599436532 | 0.014079728 |
| ***ABCA1*** | -0.068070967 | 0.859222104 | 2.207383944 | 6.43E-05 |

**Supplemental Figure 1. S1.1** Experimental steps to induce HemSC to endothelial differentiation. VEGF-B at 10ng/ml added to serum starved HemSC on Day 0 (n=6). Cells were treated ± R(+) propranolol (20 μM) for 2 hours on day 4 and day 6. **S1.2** KEGG pathway analysis indicates steroid biosynthesis and axonal guidance genes are differentially regulated in differentiating HemSC ± R(+) propranolol. **Table S1** Log$_2$ fold changes and adjusted p values of differentially regulated MVP genes and ABCA1 upon R(+) propranolol treatment on Day 4 and Day 6.

#### Supplemental Figure 2

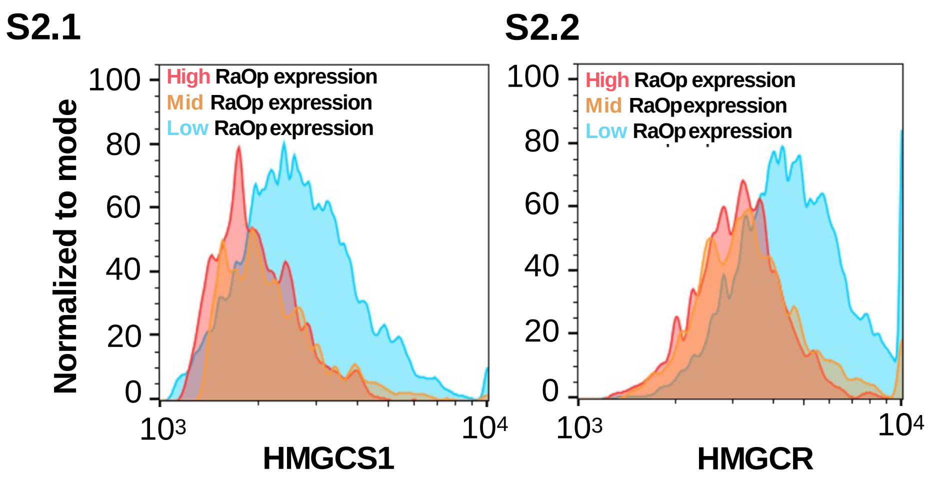


**Supplemental Figure 2.** Flow cytometry plots of HUVECs expressing fluorescently tagged SOX18$^{RaOp}$ stained for HMGCS1 (**S2.1**) and HMGCR (**S2.2**) sorted into low, intermediate, and high overexpression of *RaOP* for analysis in **Figure 2D**,**E**.

#### Supplemental Figure 3

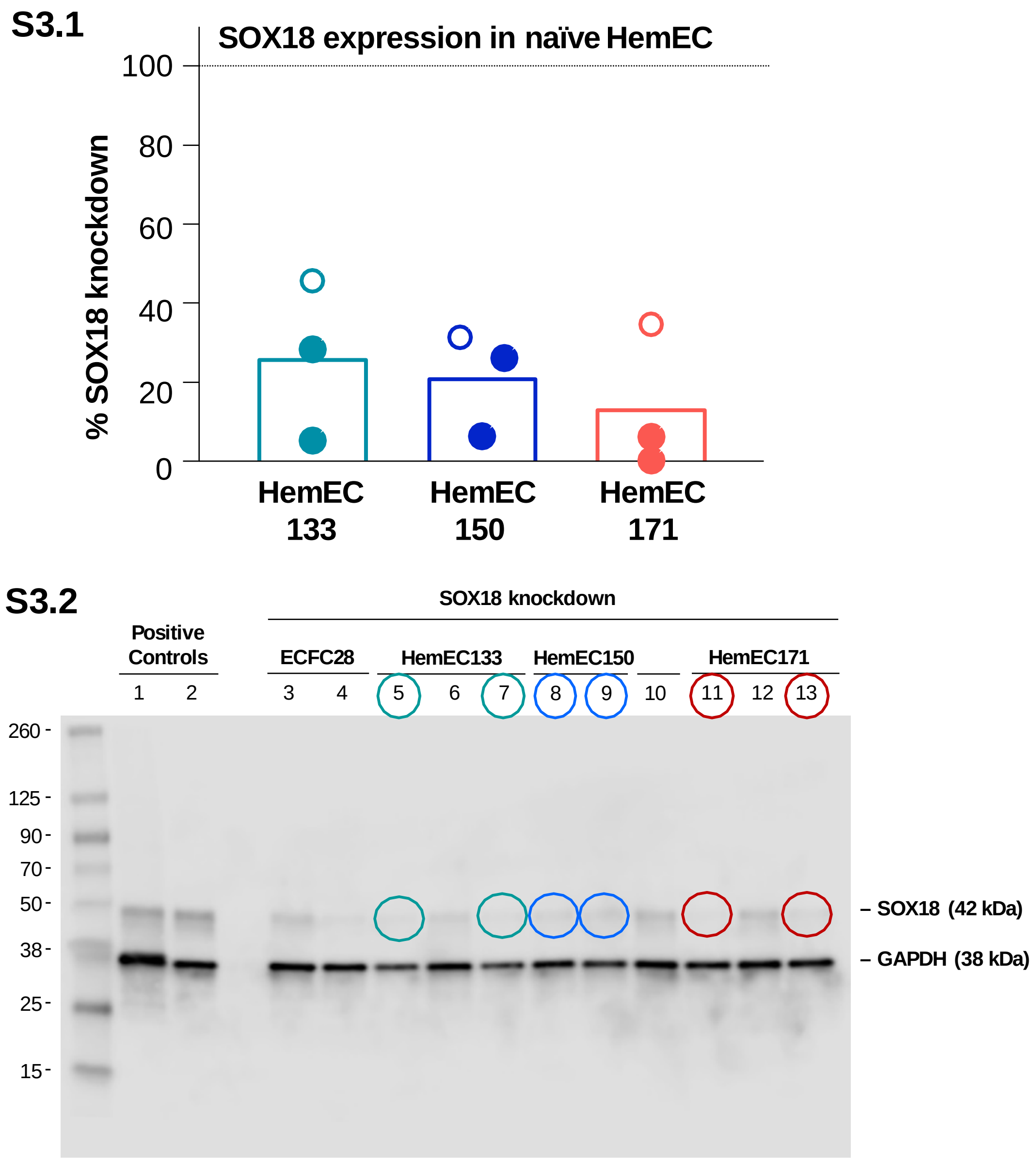


**Supplemental Figure 3**. Efficiency of lentiviral knockdown of SOX18 on mRNA (**S3.1**) and protein (**S3.2**) level measured by qPCR and Western Blot. HemEC with >70% knockdown of SOX18, shown by circles, were used for experiments in Figure 3. The lentiviral SOX18 knockdown was performed independently three times in each of the three patient-derived HemEC (denoted as HemEC 133$^{shSOX18}$, HemEC150$^{shSOX18}$, HemEC171$^{shSOX18}$) leading to a sample size of n=6 (with 3 biological replicates and 2 independent lentiviral knockdowns, with an efficacy >70% each).

### Supplemental Figure 4

#### S4.1 Skin Control (12-month-old male)

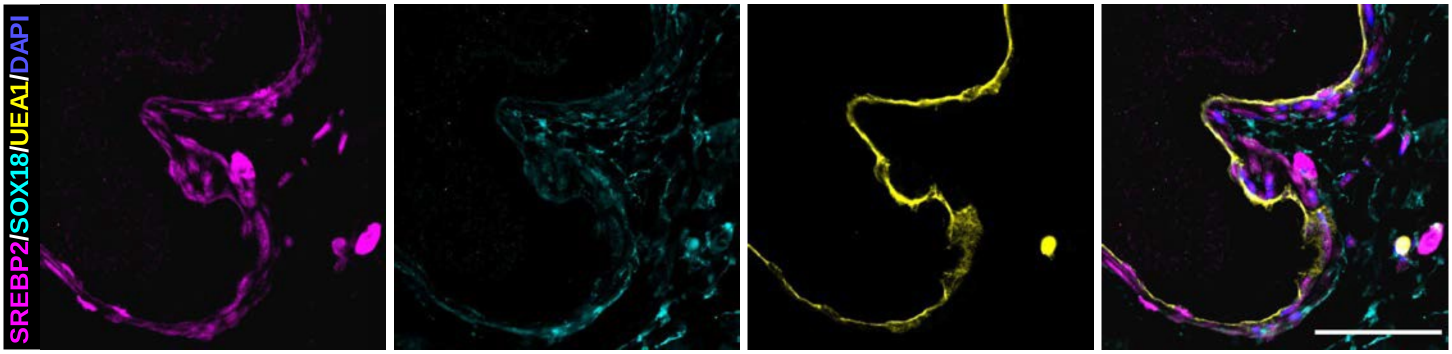


#### S4.2 Proliferating IH (7-month-old female)

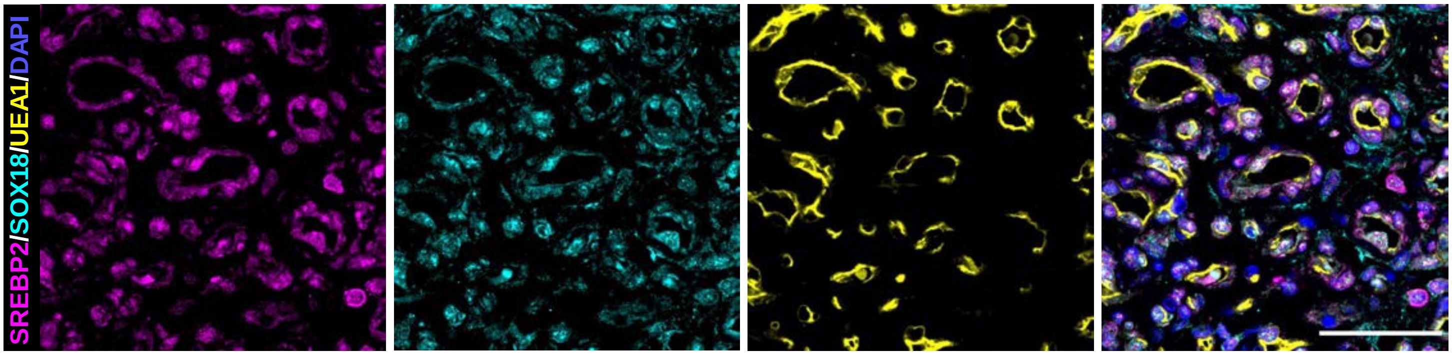


#### S4.3 Involuting IH (2-year-old female)

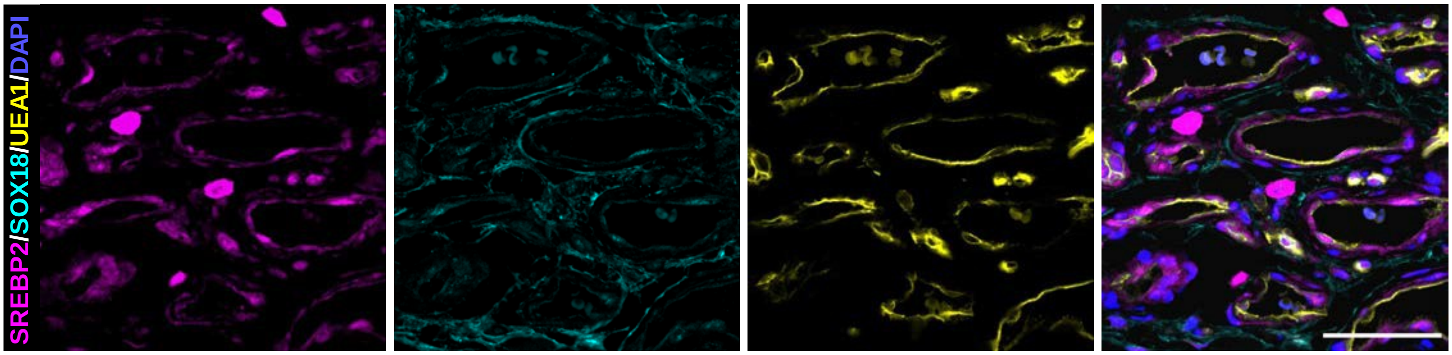


#### S4.4 Regrowing IH (6-year and 4-month-old female)

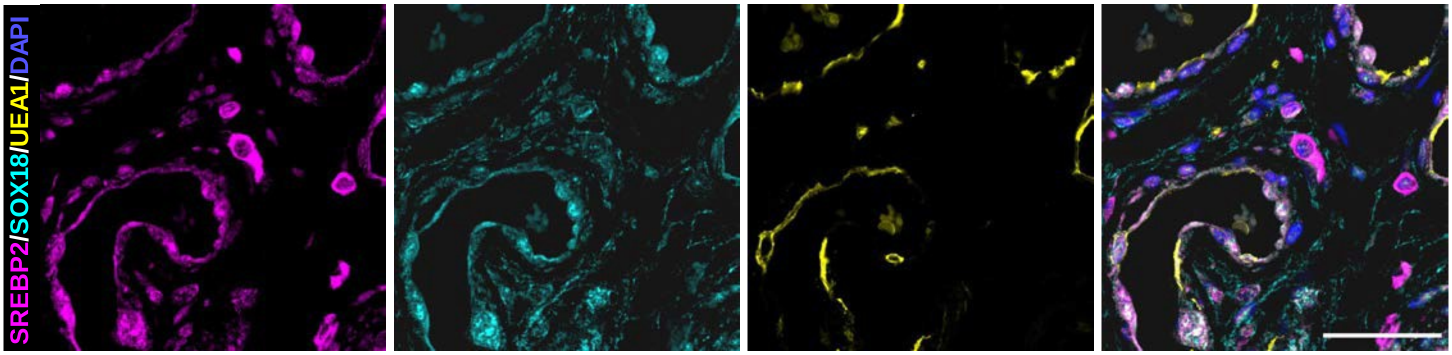

### Supplemental Figure 4, continued

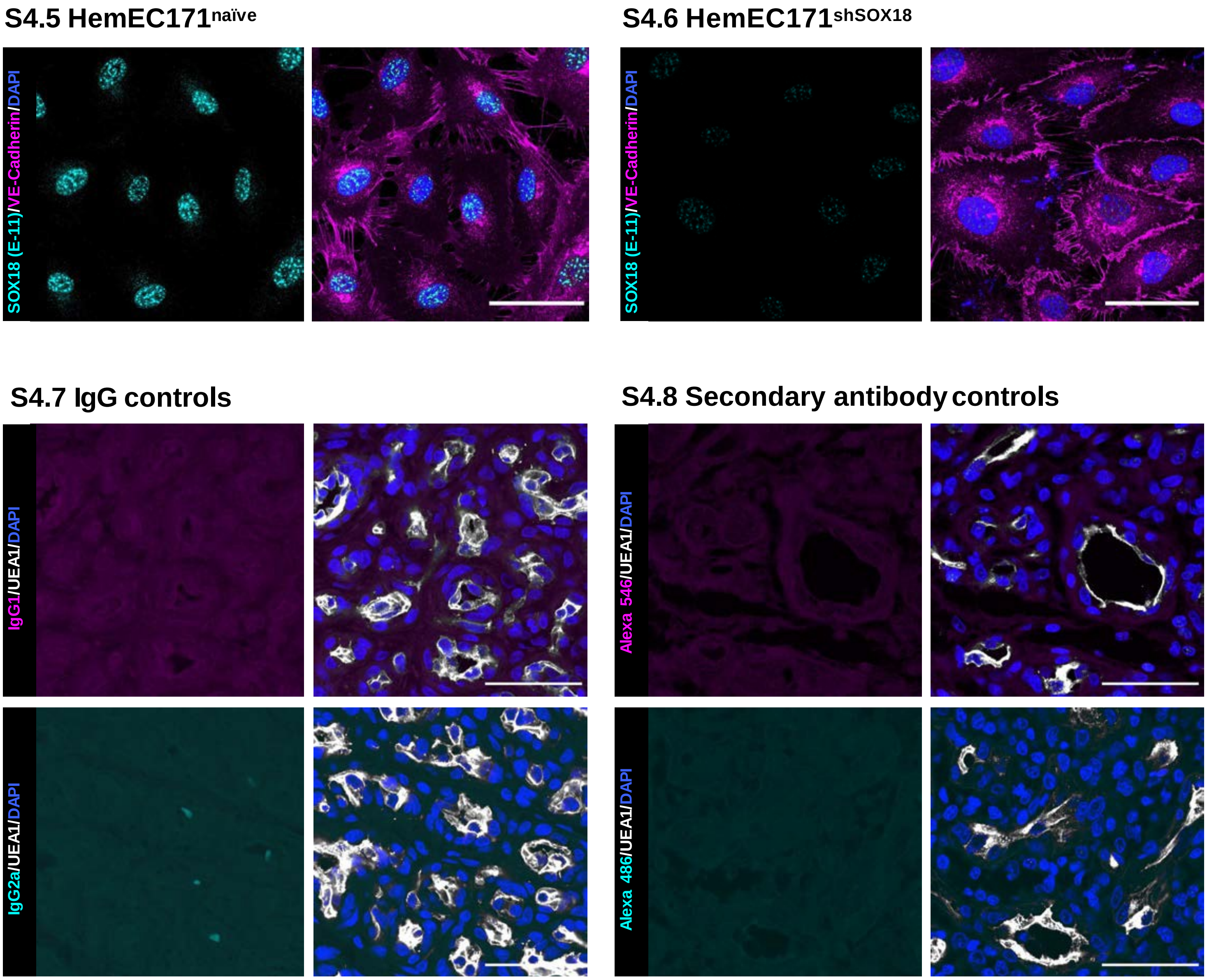


**Supplemental Figure 4. S4.1-4** Single fluorescent channels for each antibody in control skin, proliferating, involuting phase and regrowing IH stained for: SREBP2 (magenta), SOX18 (cyan), and the human specific lectin UEA1 (yellow). Cell nuclei stained with DAPI (blue). Scale bars 50 µm. **S4.5**,**4.6** SOX18 (E-11) antibody validation for specificity in HemEC$^{\text{naïve}}$ and HemEC$^{\text{shSOX18}}$ cells. (**S4.7**,**4.8**) Isotype-matched antibody as well as secondary antibody control staining of IH sections. Scale bars 25 µm.

### Supplemental Figure 5

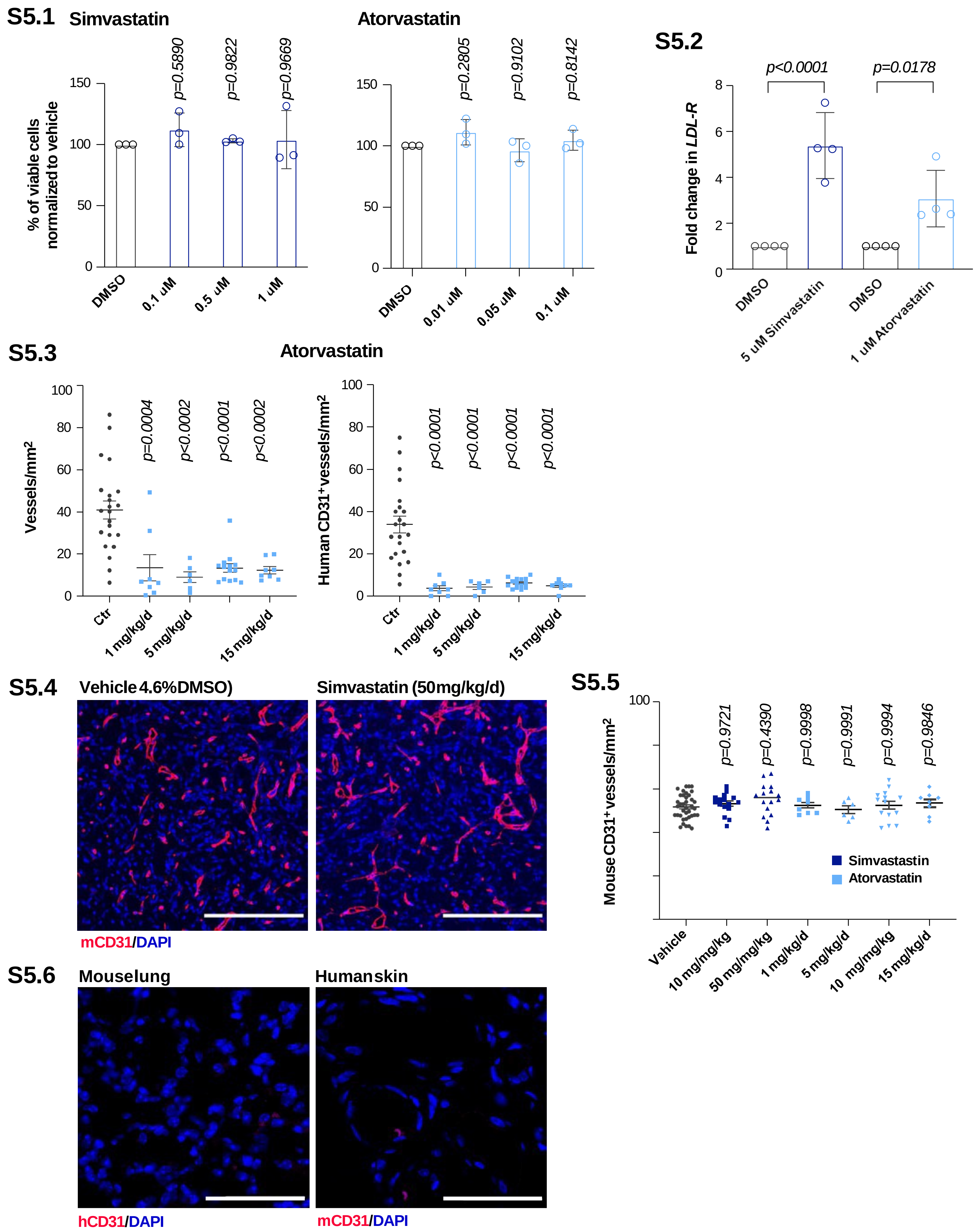

#### Supplemental Figure 5, continued

**Table S5.1: Conversion of statin doses from mouse to human**

| **Simvastatin** | | | | | | |
|---|---|---|---|---|---|---|
| Used in mice (mg/kg/d) | **50** | **10** | **5** | **1*** | **0.5** | **0.1** |
| Human equivalent dose (mg/kg/d) | 4.065 | 0.813 | 0.407 | 0.081 | 0.041 | 0.008 |
| **Atorvastatin** | | | | | | |
| Used in mice (mg/kg/d) | **15** | **10** | **5** | **1*** | | |
| Human equivalent dose (mg/kg/d) | 1.220 | 0.813 | 0.407 | 0.081 | | |

**Table S5.2 Statistic for differentiation assay, treatment with R+ propranolol, simvastatin, atorvastatin**

Two-way ANOVA with Tukey's multiple comparisons test

| | VE-Cadherin | CD31 |
|---|---|---|
| **Sample Comparison** | **Adjusted p-value** | **Adjusted p-value** |
| DMSO vs. *DMSO | 0.0693 | 0.0748 |
| DMSO vs. R(+) Prop | 0.9477 | 0.3086 |
| DMSO vs. Simvastatin | 0.1467 | 0.2395 |
| DMSO vs. Atorvastatin | 0.0155 | 0.5291 |
| *DMSO vs. R(+) Prop | <0.0001 | 0.0407 |
| *DMSO vs. Simvastatin | 0.0017 | 0.0018 |
| *DMSO vs. Atorvastatin | 0.0180 | 0.0523 |
| R(+) Prop vs. Simvastatin | 0.0183 | 0.3652 |
| R(+) Prop vs. Atorvastatin | 0.1373 | 0.8783 |
| Simvastatin vs. Atorvastatin | >0.9999 | 0.6882 |

**Supplemental Figure 5. S5.1** Cell viability of statin-treated differentiating HemSC. Simvastatin (0.1 - 1 µM) or atorvastatin (0.01 - 0.1 µM) had no effect on cell viability over the course of endothelial differentiation for 6 days. **S5.2** Simvastatin and atorvastatin increased LDL receptor mRNA as expected due to HMGCR inhibition. **S5.3** Patient-derived HemSC (n=4) were pretreated with 0.5 µM atorvastatin or vehicle (0.005% DMSO) for 24 hours, suspended in Matrigel with 0.25 µM atorvastatin or an equivalent DMSO concentration (0.0025%) and injected subcutaneously into nude mice with 2 implants/mouse. Mice were treated with 1, 5, 10 or 15 mg/kg/d atorvastatin or an equivalent volume of PBS with a DMSO concentration of max. 2.7% every 12 hours for 7 days. Treatment with atorvastatin leads to a significant reduction in vessel density at each dose. Vessel density is expressed in vessels/mm$^2$. **S5.4** The density of murine blood vessels in the Matrigel implants were unaffected by either maximum dose of simvastatin or atorvastatin compared to vehicle (as quantified in **S5.5**). **S5.6** Stainings of murine lung with anti-human CD31 and human skin with anti-mouse CD31 with respective human and mouse CD31 antibodies used throughout Figure 5 each show negative staining demonstrating specificity for human or mouse CD31 staining, respectively. Scale bars 100 µm (**S5.4**, **S5.6**). P values were calculated using one-way ANOVA multiple comparisons test with Dunnett-correction (**S5.1**), one-way ANOVA with Šidák-correction (**S5.2**), one-way ANOVA multiple comparisons test with Tukey-correction (**S5.3**, **S5.5**). Data show the mean ± SD and were collected for 2 implants in each mouse, leading to an observation sample size of n=22 for vehicle (combined), n=8 (1mg/kg/d), n=6 (5 mg/kg/d), n=14 (10 mg/kg/d), and n=8 (15 mg/kg/d). **Table S5.1** compares the reduction of vessel density observed at 1 mg/kg/d for simvastatin and atorvastatin (*) to the human equivalent doses of simvastatin and atorvastatin, shown below. The red box highlights the human equivalent dose of simvastatin used in infants with Smith-Lemli-Opitz syndrome (0.5 - 1 mg/kg/d). **Table S5.2**. Summary of statistical analysis of differentiating HemSC treated with R(+) propranolol, simvastatin, and atorvastatin in **Figure 5A**.

### Supplemental Table 6

#### Table S6: List of primers used throughout the study (Figure 1, 3, 5)

| *Gene* | *Forward* | *Reverse* |
|---|---|---|
| ***ATP5B*** | CCACTACCAAGAAGGGATCTATCA | GGGCAGGGTCAGTCAAGTC |
| ***SOX18*** | CAAGATGCTGGGCAAAGCGTG | GCGGGGGCGCTAATCC |
| ***HMGCS1*** | ACACAAGATGCTACACCGGG | TGGGTGTCCTCTCTGAGCTT |
| ***HMGCR*** | AGTGAGATCTGGAGGATCCAA | GATGGGAGGCCACAAAGAGG |
| ***MVK*** | AGATCCCAAACCCGCTGAAG | CCTTGATGGTATCGGAGGGC |
| ***ABCA1*** | CAGAGGTGGCTCTGATGACC | TGTTTTGCTTTGCTGACCCG |
| ***CD31*** | CACCTGGCCCAGGAGTTTC | AGTACACAGCCTTGTTGCCATGT |
| ***VE-Cadherin*** | CCTTGGGTCCTGAAGTGACCT | AGGGCCTTGCCTTCTGCAA |
| ***NOTCH1*** | CGGTGAGACCTGCCTGAATG | GCATTGTCCAGGGGTGTCAG |
